# Primate Cerebellar Scaling in Connection to the Cerebrum: A 34-Species Phylogenetic Comparative Analysis

**DOI:** 10.1101/2023.03.15.532597

**Authors:** Neville Magielse, Roberto Toro, Vanessa Steigauf, Mahta Abbaspour, Simon B. Eickhoff, Katja Heuer, Sofie L. Valk

## Abstract

The cerebellum has increasingly been recognized for its role in diverse functional processes. The reciprocally connected cerebello-cerebral system may scaffold both brain size increase and advanced associative abilities, evolving highly coordinately in primates. In parallel, functional cerebello-cerebral modules have undergone reorganization and cerebellar lobules crura I-II (the ansiform area across mammals) have been reported to be specifically expanded in humans. Here we manually segmented 63 cerebella (34 primate species; 9 infraorders) and 30 crura I-II (13 species; 8 infraorders). We show that both constraints and reorganization may shape the evolution of the primate cerebello-cerebral system. Using phylogenetic generalized least squares, we find that the cerebellum scales isometrically with the cerebral cortex, whereas crura I-II scale hyper-allometrically versus both. Our phylogenetic analyses evidence primate-general crura I-II hyperscaling in contrast to virtually isometric cerebello-cerebral scaling. Crura I-II hyperscaling may be important for associative and cognitive brain functions in an evolutionary context.

## Introduction

The cerebellum is implicated in abstract functions such as executive, social, and emotional processing^1–4^. Lesion studies^5,6^ evidence behavioral deficits in these functional domains. The stereotyped structural organization of the cerebello-cerebral system, which has undergone marked modular reorganization in primates, likely supports its diverse functions^7–10^. Human cerebellar involvement in cognitive modules^11–13^ may explain its activation in diverse cognitive tasks^14,15^ and its role in functional connectivity networks^16–18^.

The cerebellum contains multiple output channels^11,12,19,20^, which are organized into multiple reciprocal loops with the cerebral cortex^12,21–24^. It likely forms internal models encoding mental representations from separate streams, returning updated predictions^25^. Cerebellar structural stereotypy and phylogenetic conservation^26^ argue for algorithmic^27,28^ uniformity across functional domains^24,29–31^, and a direct link between function and cerebello-cerebral connectivity^7,12^. Distinct cerebellar computations may still occur^29^ and integration cannot be excluded based on overlapping connections^32^, fractured somatotopy^33^, and transmodal integration in single granule cells^34^.

Similar to humans^21,22^, non-human primate (NHP) hemispheric cerebellar lobule VII, and especially crura I-II, connect reciprocally to prefrontal cortex (PFC)^11,12,23,35^. A cross-species crura I-II homolog has been referred to as the ansiform area^36,37^. We adopt this term here. Functionally, cerebellum-PFC connectivity modulates cognitive processing speed^38^, congruent with reciprocal functional coherence between PFC and ansiform area^39^. Human crura I-II additionally connect reciprocally to temporal and parietal associative cortices^21,22^. Resting state fMRI (rs-fMRI) studies illustrate widespread ansiform area involvement in functional networks, with extensive functional connectivity to parietal cortices including Brodmann area 7^40,41^. Analyses ranging from winner-takes-all parcellations^16,17,42^, to gradients^43^ and lead-lag relationships^42^ indicate its unique involvement in associative networks such as the default mode (DMN), control, and frontoparietal networks, and its extreme position along the functional gradient separating DMN and language networks from motor functions. In humans, consistent with dominant contralateral cerebello-cerebral connectivity^21,22^ and conserved functional organizational asymmetry between macaques and humans^44^, meta-analyses show left-lateralized executive and right-lateralized language ansiform area activations^14^.

Primate brain size increase^45–47^ comes primarily from expansion of the cerebello-cerebral system^48^. This system’s structure and connectivity enable diverse functional specializations^7,10,49^. Primate cerebellar and cerebral volumes^50,51^ and neuron numbers^52,53^ increase highly coordinately. Other vertebrates with abstract reasoning abilities^54–56^ have similar cerebello-cerebral anatomy and scaling^57,58^ as primates^57^, suggesting strong conservation across phyla^59^. Meanwhile, primate evolution seems to also involve reorganization of cerebello-cerebral areas and networks, exceeding volumetric increases^60^ in their importance for cognitive abilities^46,47,61^.

Previous work has revealed phylogenetic axes of functional organization^62,63^. Structural modularity^64,65^ within the primate cerebello-cerebral system may reflect functionally relevant reorganization with functionally and structurally connected subregions evolving at increased relative pace. Such disproportionate expansions of functionally-related structures may indicate behaviorally relevant changes, as they persist over millions of years^66^ and against great metabolic costs^67–69^. In great apes, cerebellar volumetric expansion may have outpaced that of the cerebrum^70^. Similarly, cerebellar surface area expansion may have outpaced that of the cerebral cortex between macaque monkeys (33% of the cerebral cortex) and humans (78%)^71^. Relatively large lateral cerebellar hemispheres characterize four vertebrate clades including primates^9^ and especially hominoids^72^. Specifically, the ansiform area volume fraction of the cerebellum is larger in humans than in macaques^13^, and appears generally larger in most primates than in mice and rats^36^. Expansion of lateral cerebellar areas alongside cerebral association areas in primates^7,73–75^ putatively reflects selection on cerebello-cerebral networks^8,9,13,50,76^

In the current study, we aimed to expand comparative analyses of the primate cerebello-cerebral system. We used a large (34 species, 63 specimens; Figure 1) MRI dataset alongside phylogenetic comparative methods to evidence isometric scaling between the cerebellum and cerebrum^50,51,77^ and assess ansiform area hyperscaling^13,36^. Where possible, we explored intraspecific variability, which is substantial among brain traits^72,78–82^ and an inherent challenge for comparative primatology^83^.

**Figure 1:**
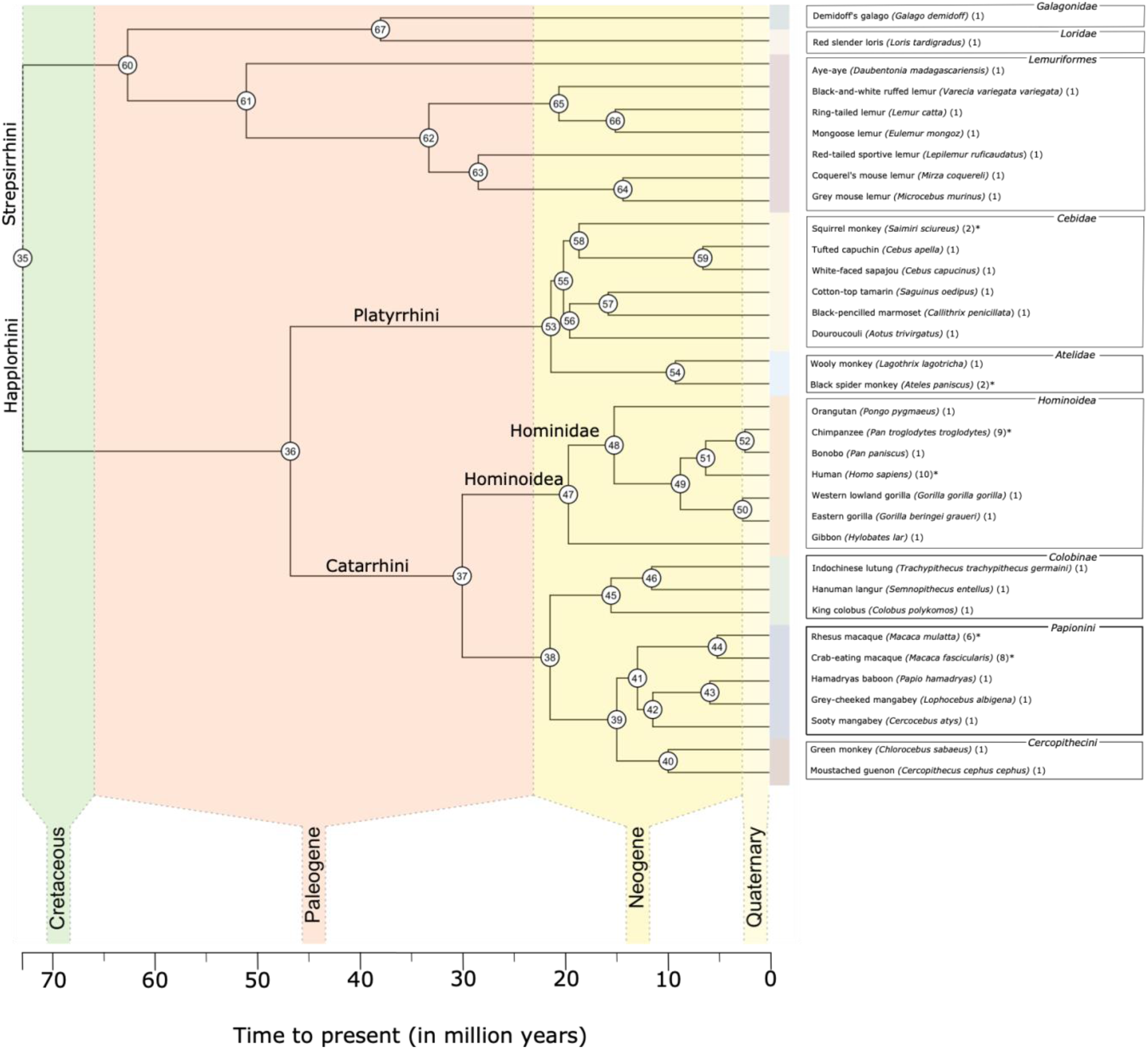
Consensus phylogenetic tree for the 34 primate species in this study. We obtained the consensus tree for the 34 species in our dataset from 10kTrees **Arnold et al. (2010)**. Archeological time periods are superimposed over the tree to provide a temporal perspective of predicted species bifurcations. Internal node numbers are plotted and can be used to identify ancestral characters (**Supplemental Table S1**). Extant species included in the current study are given on the right and colored by their clade membership. Additionally, the numbers of specimens per species are given, and species with multiple specimens are marked with asterisks.

Our analyses confirmed isometric scaling between the cerebellum and cerebrum. Reanalysis of Stephan collection^84^ data indicated this relation may be hypo-allometric. Conversely, the ansiform area scaled hyper-allometrically versus the cerebellum and cerebrum. Contrary to our hypothesis based on previous work^13,70,72^, hominoids did not exhibit exceptional scaling.

## Methods

### Overview

We perform phylogenetic comparative analyses on manually segmented cerebella from 34 primate species (N=63; Figure 1) and ansiform areas from 13 species (N=30). We provide here an overview of the methods. For a more detailed description see Supplemental Methods and for the full protocol see Heuer et al. (2019)^85^.

### Data

Primate comparative data are rare and sizable datasets are difficult to obtain. We used the collection of primate species collated from different sources in the Brain Catalogue Primates project^85^, available on BrainBox^83^. It included 35 species (N=66), which covered the primate phylogenetic tree relatively well, with 9 infraorders from Strepsirrhini and Haplorhini suborders. The red howler monkey was excluded due to irreparable damage. Cerebellar damage was also noted in other specimens, whose volumetric estimates need to be treated with due diligence. Cebidae (7 species), Hominoidea (7), Lemuriformes (6), and Papionini (5) were most extensively sampled (Figure 1). Scans included both fully abstracted brains and *in situ* brains, either full body scans or only including the skull. T1- or T2-weighted (T1w or T2w) MRI anatomical scans were obtained at either 1.5, 3T, 7T or 11.4T for up to 12 hours. See Supplemental Methods for basic data acquisition, provenance^85–89^ and damage description, and Supplemental File S1 for quality-control data including signal-to-noise ratios and resolutions.

### Manual segmentation procedure

Manual segmentation provides ground truth for brain structure volumes^90^. We obtained initial cerebellar masks semi-automatically, subtracting cerebral masks from whole brain masks with StereotaxicRAMON (github.com/neuroanatomy/StereotaxicRAMON) or using Thresholdmann (github.com/neuroanatomy/thresholdmann) to interpolate between thresholds at manually selected locations. Initial cerebellar masks were uploaded to BrainBox^91^ for online collaborative segmentation of MRI images. Lastly, we used StereotaxicRAMON for mathematical morphology operations that preserve original mask topology (Figure 2a).

**Figure 2:**
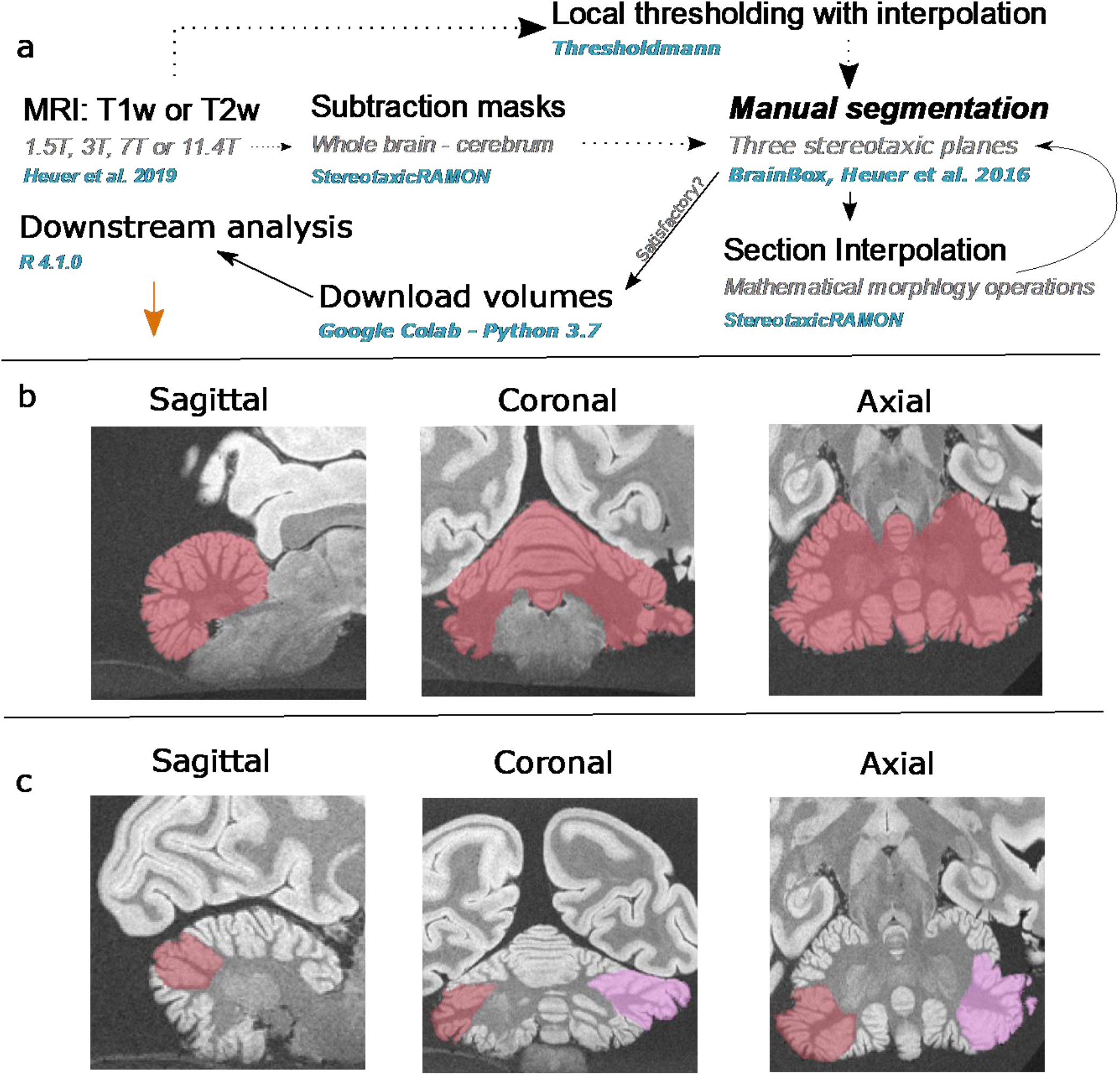
Manual segmentation method for primate brains. **(a)** A schematic representation of the manual segmentation pipeline. T1- and T2-weighed (T1w and T2w) MRI brain scans previously described in **Heuer et al. (2019)** were used to make initial cerebellar masks, by subtracting cerebral masks from whole brain masks with StereotaxicRAMON. Some initial cerebellar masks were made through local thresholding and interpolation with Thresholdmann. Cerebellar masks were then uploaded to BrainBox, where manual segmentation was performed. Manual segmentation was performed iteratively with interpolation between slices with mathematical morphology operations, until segmentations reached satisfactory quality. Volumes were then downloaded with a custom python script for subsequent analysis in R. **(b-c)** Example segmentations in stereotaxic planes for the cerebellum and ansiform area in the hamadryas baboon. **(b)** Cerebellar segmentations primarily involved removal of the brain stem, removing erroneously marked tissue and sulci, and reconstructing damaged tissue. **(c)** Ansiform area segmentations were slightly more challenging and additionally involved identification of superior posterior and ansoparamedian fissures, and segmentation between them. Segmentations were made so they did not enter vermal portions of lobule VII.

### Cerebellum

Manual segmentations (Figure 2b) consisted of brainstem removal, cerebellar surface boundary determination, erroneously marked sulci erasure, and damaged tissue reconstruction. Segmentations were performed by NM, with SLV providing preliminary segmentations for a subset of specimens. Damaged tissue was reconstructed by carefully manually interpolating between non-damaged slices. Segmentations were performed to satisfaction using the three stereotaxic planes and alternating the manual segmentation procedure and mathematical morphology operations. All segmentations (N=65) were approved by KH and SLV.

### Ansiform area

Segmentation (Figure 2c) relied on identification of superior posterior (SPF) and ansoparamedian (APMF) fissures^13,92,93^. Since data quality was variable, segmentation was performed in a subset of specimens representative of the sample that allowed fissure identification with reasonable certainty: 5 apes (N=22; including 10 humans and 9 chimpanzees) and 8 non-apes. Segmentations were again iteratively performed to satisfaction. An interobserver strategy validated segmentations made by NM. For six species, a secondary observer (MA) provided blinded segmentations alongside NM. Additionally, a tertiary observer (VS) segmented the remaining human and chimpanzee ansiform areas in consultation with NM. All segmentations were checked by KH and SLV.

### Reliability of segmentations

Reliability of ansiform area segmentations was assessed by ANOVA:

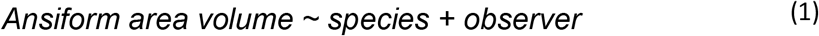

Human cerebellar and ansiform area manual segmentations were compared with automated cerebellar and lobular segmentations made using CERES^94^ at volbrain.upv.es:

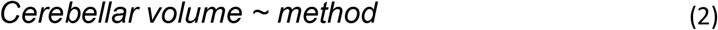

and

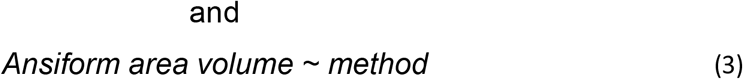

Intraclass correlation (ICC) was additionally calculated^95^. ICC of .75-.90 indicates good reliability and .90-1.00 excellent reliability^96^.

Cerebellar volume was significantly related to method (Pr(>F)<.01). ICC was .26 (95% confidence interval (CI): -.38<ICC<.74). Inspection revealed that manual segmentations were on average 20.4 cm^3^ (16.2%) larger than CERES segmentations. Ansiform area volume was not related to method (Pr(>F)=.286), with ICC of .84 (95%CI: .51<ICC<.96). Manual volumes were only 2.7 cm^3^ (6.6%) larger. One specimen differed by 6.9 cm^3^ (18.2%). It represented an average-sized cerebellum, unlikely to alter the results significantly. Full results can be found in the GitHub repository.

Manual ansiform area volumes were significantly related to species (Pr(>F)<.001). No significant observer effect was detected (Pr(>F)=.468). ICC was .99 (95%CI: .92<ICC<1.00).

### Neuroanatomical measurements

Phenotypes of interest included cerebellar, cerebral^85^, and ansiform area volumes, and cerebellum-to-cerebrum and ansiform area-to-cerebellum ratios. Absolute measurements were log10-transformed. Volumes were not corrected for shrinkage. Normality was examined for traits with multiple (>2) observations per species (Figure 1) by Shapiro-Wilk tests^97^ and outliers were visualized on boxplots (±1.5 IQR from Q1 and Q3). This led to the exclusion from subsequent analyses of one rhesus and one crab-eating macaque (Supplemental Figure S1). Two different crab-eating and rhesus macaque specimens (Supplemental Figure S2) displayed cerebellum-to-cerebrum ratio outliers but were retained in analyses as cerebellar and cerebral volumes fell within the normal range.

### Statistical analyses and reproducibility

Consensus phylogenetic trees for the 34- (Figure 1) and 13-species analyses were obtained from the 10kTrees^66^ primate database (10ktrees.nunn-lab.org; version 3). They were constructed from 17 genes and 7 different loci^66^ (10ktrees.nunn-lab.org/downloads/10kTrees Documentation.pdf).

We used R version R 4.1.0 for statistical analyses^98^. Fit of extant phenotypes in the current and previous^85^ studies were tested with three common evolutionary models of trait evolution using Rphylopars^99^. Models included Brownian Motion (BM)^100^, Ornstein-Uhlenbeck (OU)^101^ (with single alpha, alpha per phenotype, or full multivariate alpha matrix), and Early Burst (EB)^102^. BM describes traits varying randomly over time in direction and extent, leading to differences predictable by time since divergence^103^. BM does not equal genetic drift but may represent weak selection within BM parameters or selection towards specific trait optima whose distribution is described by BM^104^. OU processes behave like BM with a specific trait optimum, a phenotype with adaptive advantage^105,106^. Lastly, EB processes^102^ – rapid change after adaptive radiations followed by comparative phenotypic *stasis* – may explain trait evolution. To assess phylogenetic signal within the data, we chi-squared tested BM with Pagel’s λ=1.0 (full phylogenetic signal) versus λ=0.0 (star-model, no phylogenetic signal). Data fit was determined by Akaike information criterion (AIC)^99,107^, with lower values representing better fit. A difference of four to seven points represents significantly less support and a difference exceeding ten indicates virtually no support for the lower-performing model.

Ancestral character estimations (ACEs) were constructed by mapping extant trait values back in time following the best-supported evolutionary model, incorporating intraspecific variation. ACEs were mapped to consensus phylogenetic trees for visualization.

Extant primate traits are not statistically independent, but share variable amounts of evolutionary history^103^. Substantial phylogenetic signal was detected, necessitating correction. Corroboratively, correlations between current and previously^85^ recorded phenotypes illustrated severe collinearity (Figure 3). Hence, we performed regressions with phylogenetic generalized least squares (PGLS)^103,108^ implemented in *nlme*^109^, which under BM^100^ generalizes to phylogenetic independent contrast (PIC) regression forced through the origin^110^. PICs^103^ were calculated with ape’s^111^ *pic.ortho* function, accounting for intraspecific variation. Regressions were performed for cerebellar volume regressed on cerebral volume for all species (1) and those with complete (including ansiform area) data (2), and ansiform area volume regressed on rest of cerebellar volume (3) and cerebral volume (4), all taking species medians. ANOVA assessed if apes and non-apes showed different scaling in all regressions. We calculated R^2^_likelihood_^112,113^ for full regressions, and apes or non-apes separately. Fisher’s R-to-Z transformation (cran.r-project.org/web/packages/psych/index.html) determined improvement of fit.

**Figure 3:**
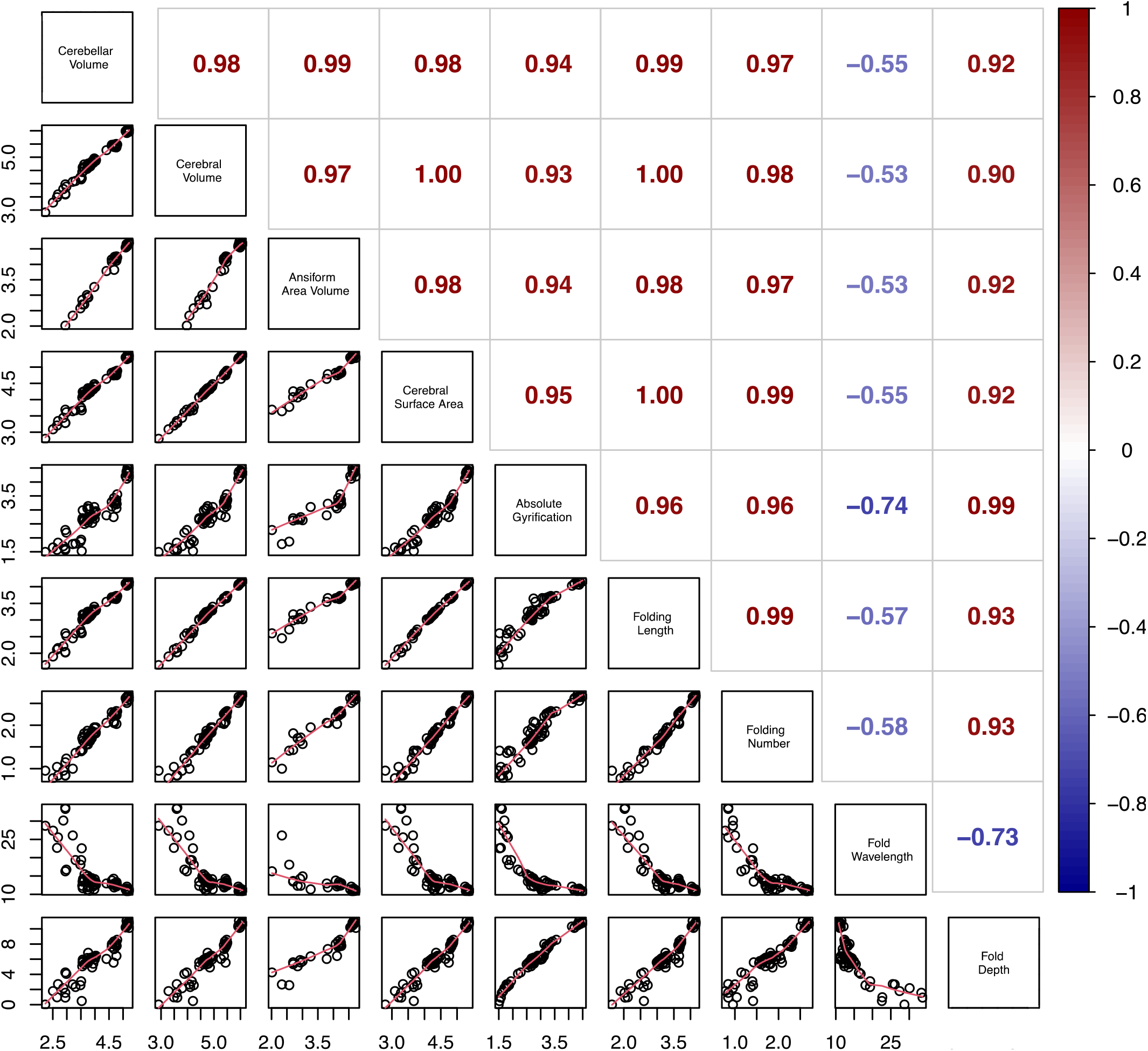
Correlations for neuroanatomical measurements. Volumes recorded in the current study were correlated with the measurements from **Heuer et al. (2019)**. The lower diagonal displays the correlations of log_10_-transformed variables, which were not corrected for phylogeny and included all specimens. These correlations serve to illustrate the strong collinearity between the neuroanatomical variables, which are an expected outcome of allometric scaling. Corresponding R^2^ coefficients for the correlations can be found on the upper diagonal. Cerebellar, cerebral, and ansiform area volumes correlated strongly and positively with all other variables, except fold wavelength, for which a negative correlation was observed.

Lastly, we repeated analyses in the Stephan collection^84^. Cerebellar and cerebral volumes were matched to primates in 10kTrees^66^. This led to a partially overlapping (in species, not specimens) dataset of 34 species.

## Results

### Neuroanatomical measures

Median neuroanatomical measures are provided in Table 1. Intraspecific data spread is given as median absolute deviations (MADs). Cerebella and ansiform areas were largest in apes, with human volumes being two to three times larger than in any other species. Cerebellum-to-cerebrum ratios were relatively consistent within clades, but variable across them. They were comparatively high in lemurs and apes. Chimpanzee cerebella occupied a range of ratios comparable to other apes (median=18.36%; MAD=3.56), whereas human ratios were relatively low (median=14.26%; MAD=1.75). Ansiform area-to-cerebellum ratios were highest in great apes. Humans and chimpanzees displayed the highest ratios (30.76% and 27.25%). Several species, including the tufted capuchin (*Cebus apella*), hamadryas baboon (*Papio hamadryas*), and green monkey (*Chlorocebus sabaeus*) had high ansiform area-to-cerebellum ratios relative to cerebellum-to-cerebrum ratios. Other species including the aye aye (*Daubentonia madagascariensis*) and bonobo (*Pan paniscus*) showed the opposite relationship, suggesting that scaling of these traits may be partly dissociated.

**Table 1:**
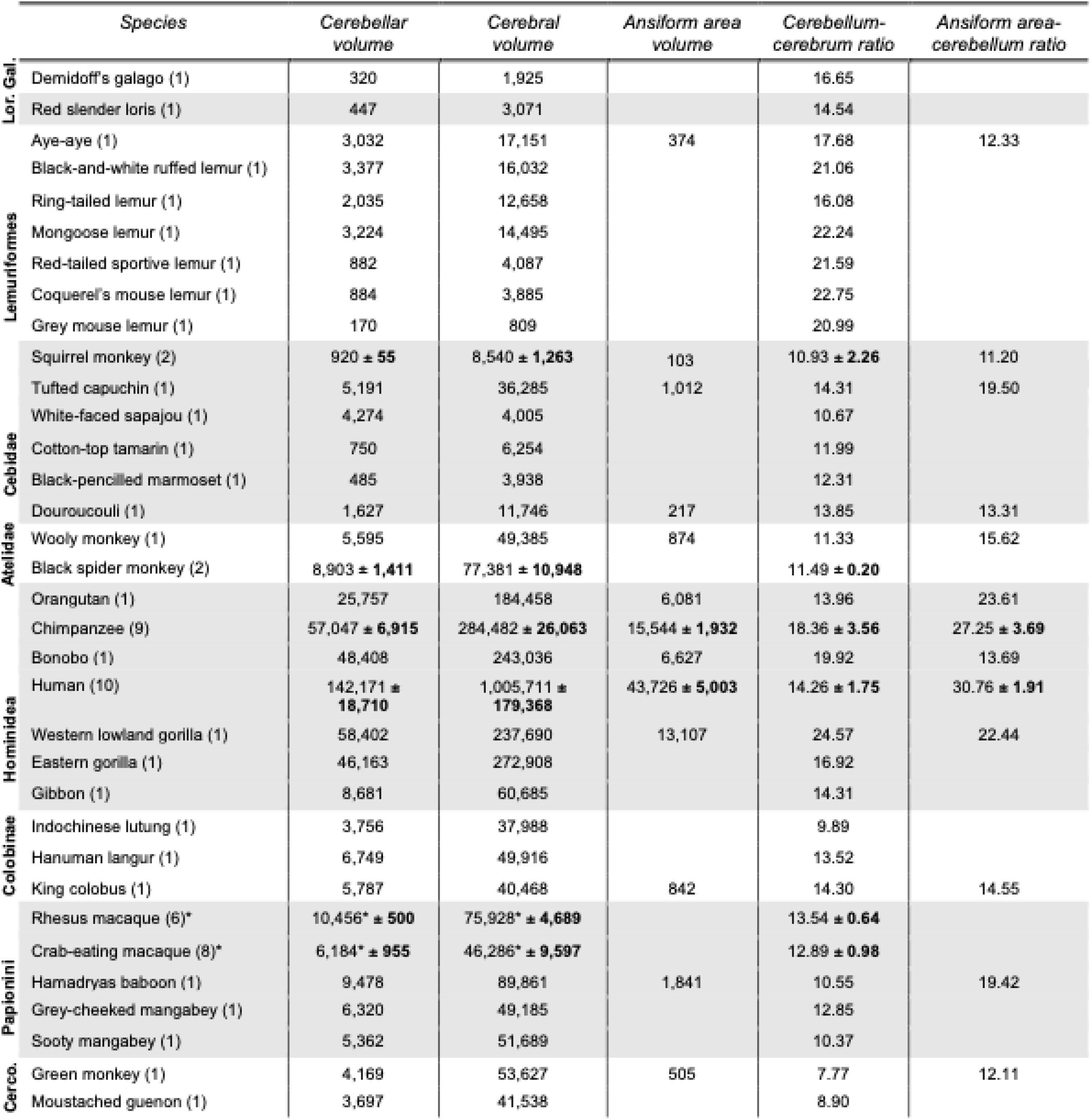
Neuroanatomical measurements. Species median measurements for previously reported cerebral volumes **Heuer et al. (2019)** are reported alongside cerebellar and ansiform area volumes (in mm^3^). Species ratios between median cerebellar and cerebral volumes, and ansiform area and cerebellar volumes are also given (in percentages). Species are ordered by the phylogenetic tree. For six species (*Ateles paniscus, Homo sapiens, Macaca fascicularis, Macaca mulatta, Pan troglodytes*, and *Saimiri sciureus*) several specimens were available. For these species, median absolute deviations are also reported. Some specimens were recorded as outliers and removed before subsequent analyses. These include a crab-eating and rhesus macaque, which were outliers in their cerebellar and cerebral volumes, as marked by the asterisks. Gal. = Galagonidae; Lor. = Loridae; Cerco. = Cercopithecini.

#### Intraspecies variability is substantial

For several species, we analyzed intraspecific variability. Variability, quantified as median absolute deviations (MAD) as percentage of the median, ranged from 4.8 to 15.8% for cerebellar volume, and from 6.2 to 20.7% for cerebral volume. Intraspecific ansiform area observations were only available for humans (median:43.73±5.00 cm^3^; MAD=11.4%) and chimpanzees (median:15.54±1.93 cm^3^; 12.4%). Intraspecific variation in cerebellum-to-cerebrum ratio ranged from 1.7 to 20.7% of the median. Range was less substantial for ansiform area-to-cerebellum ratios for humans (median:30.76±1.91%; 6.2%) than for chimpanzees (median:27.25±3.69%; 13.5%). Volume and ratio varied relatively independently across species. Ordered bar graphs for phenotypes can be found in Supplemental Figure S3.

### Evolutionary model testing

AIC ranking^114^ revealed the following support for evolutionary models: Brownian Motion (BM) (λ=1.0; AIC=6209.62) > Early Burst (EB) (AIC=6216.25) >> BM (λ=0.0 (star phylogeny); AIC=6267.87) >> Ornstein-Uhlenbeck (OU) (single α; AIC=6609.29) >> OU (α per phenotype; AIC=7143.13) >> OU (full multivariate α; AIC=8063.37). The BM was best-supported, and significantly more so than EB. Chi-squared testing revealed strong support for the λ=1.0 versus the λ=0.0 model (χ^2^=53.68). We adopted BM for subsequent analyses.

### Ancestral character estimations

We provided ancestral volumes for cerebellar, cerebral, and ansiform area volumes using BM (Supplemental Table S1). Full ancestral character estimations (ACEs) including those reported for neocortical measures^85^ can be found in the GitHub repository.

#### Relative cerebello-cerebral traits differ across clades

We mapped ACEs to the phylogenetic tree, illustrating evolutionary dynamics of the cerebello-cerebral system. We plotted 95% confidence intervals for ACEs onto the trees (Figure 4), with uncertainty intuitively increasing with time to present (Supplemental Figure S4). Cerebellar and cerebral volumes showed virtually identical evolutionary dynamics (Figure 4a,b), with largest volumes in apes. Cerebellar volume at the ancestral node of the 34-species tree was estimated at 1856mm^3^, resembling the ring-tailed lemur in our data. When we mapped the cerebello-cerebral ratio (4c), relative cerebellar volume was increased in several clades specifically: Hominoidea, and Strepsirrhini clades including Galagonidae, Loridae, and Lemuriformes (Figure 1).

**Figure 4:**
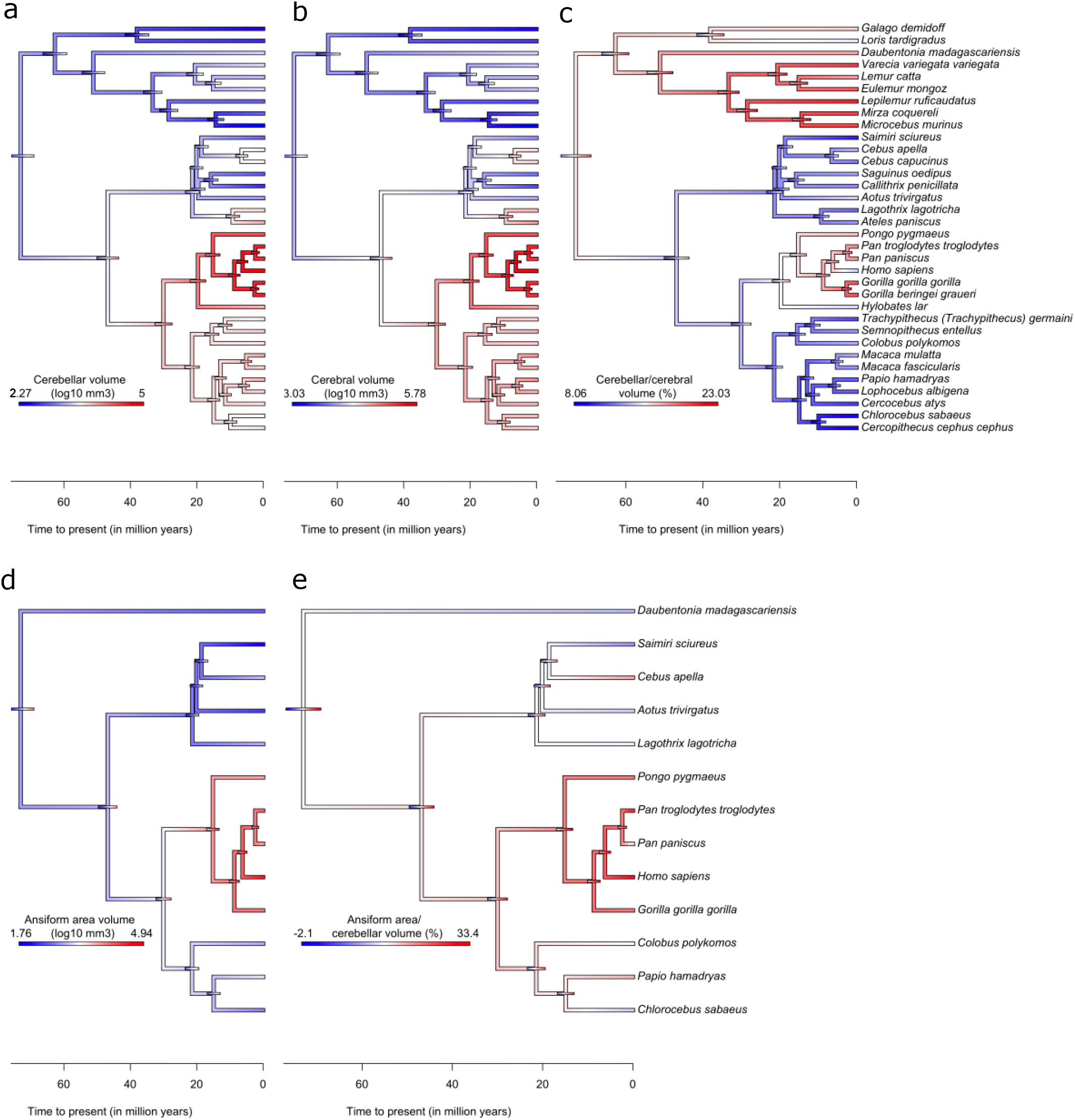
Ancestral character estimations for neuroanatomical phenotypes. Ancestral character estimations (ACEs) based on the Brownian Motion model of trait evolution are provided for absolute **(a, b, d)** and relative **(c, e)** volumes, alongside nodewise confidence intervals (colored bars on tree nodes). ACEs for cerebellar **(a)**, cerebral **(b)**, and relative cerebellar-to-cerebral **(c)** volumes were calculated from and mapped to the full 34-species tree, whereas ansiform area **(d)** and relative ansiform area-to-cerebellar **(e)** volumes were calculated from and mapped to the 13-species tree. The ancestral node was estimated at 73 million years to present.

Similarly, ansiform area volumes (4d) showed largest values in great apes as expected due to larger body size. Ansiform area-to-cerebellum ratios mapped to the tree (4e) were more interesting. In contrast to cerebellum-to-cerebrum ratios (4c), ansiform area-to-cerebellum ratios were highest in great apes, but not in the one Strepsirrhini species we measured (a lemur: aye aye; *Daubentonia madagascariensis*). Within great apes, humans (30.76%) and chimpanzees (27.25%) had the highest ratios. The tufted capuchin (*Cebus apella*; 19.50%) showed a high ratio for its phylogenetic position (versus the squirrel monkey; *Saimiri sciureus*; 10.93%+2.26). For all ancestral character estimations, we provide additional visualization in Supplemental Figure S5, which highlights the relationship between trait values and evolutionary time more directly.

### Allometric scaling relationships

#### Accelerated ansiform area scaling in primates

We employed PGLS analysis in the context of BM evolution (Figure 5). Median cerebellar volumes were regressed on median cerebral volume for full 34-species data (5a) and for the 13 species with complete data (5b). The scaling relationships were slightly below 1:1 in both cases (slope_34_=.955 and slope_13_=.940). Although trending towards hypo-allometry, cerebellar and cerebral volumes evolved isometrically. Ansiform area volume scaled hyper-allometrically to rest of cerebellar (ROC; 5c) and cerebral (5d) volumes (slope_ROC_=1.297; slope_cerebrum_=1.245), with lower intercept for cerebrum (intercept_cerebrum_=-2.784) than for ROC (intercept_ROC_=-1.833). Together, the ansiform area scaled hyper-allometrically to both ROC and cerebral volume.

**Figure 5:**
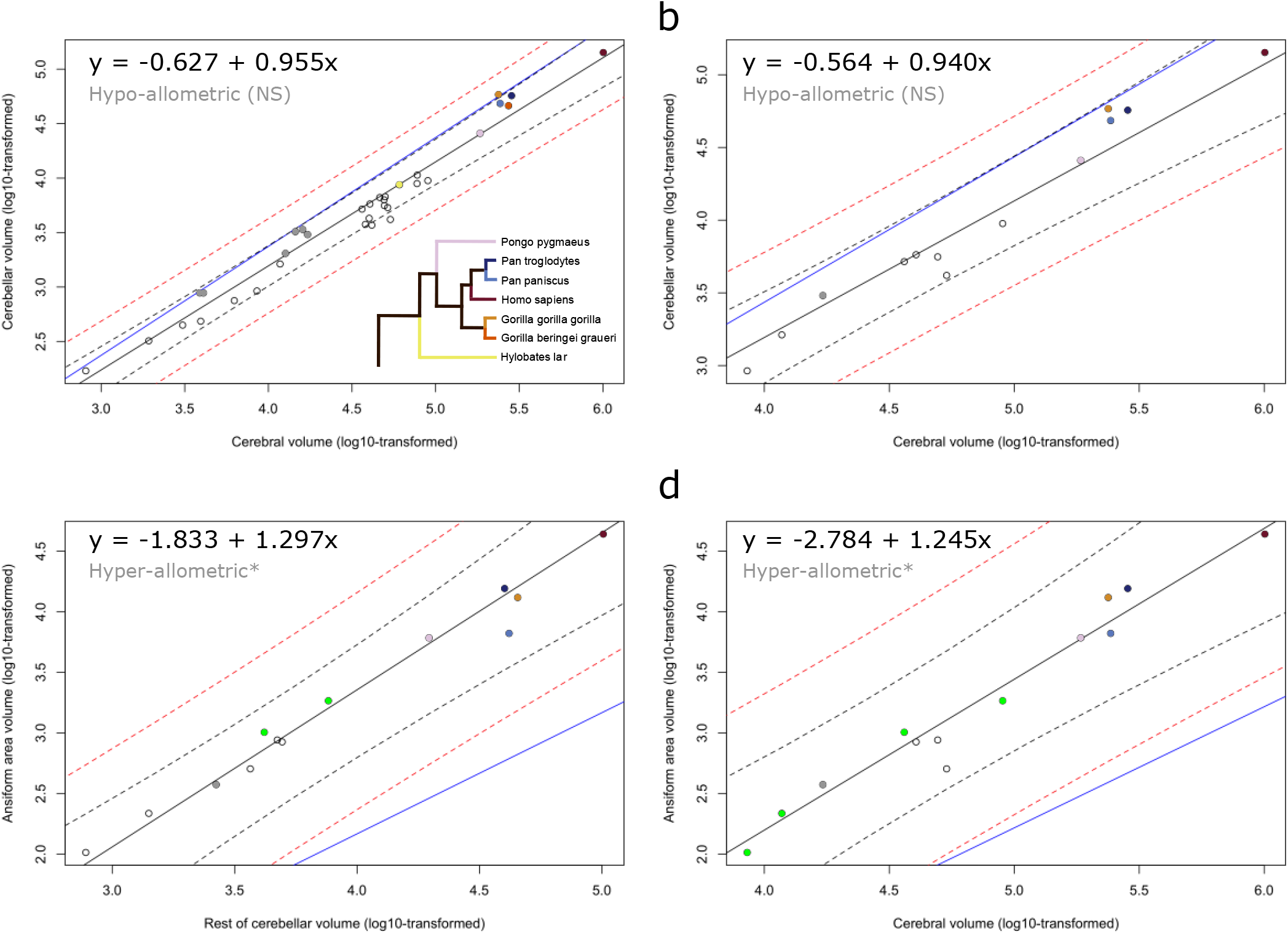
Allometric scaling relationships of cerebellar and ansiform area volumes. Phylogenetic generalized least squares (PGLS) regressions for cerebellar volume regressed on cerebral volume **(a,b)**, and ansiform area regressed on rest of cerebellar and cerebral volumes **(c,d)**. Median volumetric data were taken from 34-species **(a)** and 13-species **(b-d)** data. Log_10_-transformed neuroanatomical measures were plotted and overlaid with regression lines obtained in respective PGLS models. 95% confidence (black dotted line) and prediction (red dotted line) intervals are provided alongside a fictive isometric scaling relationship with the same intercept (blue line). Exclusion of the isometric scaling relationship from the confidence intervals was taken to indicate significant allometry, as indicated by the asterisks. **(a,b)** Cerebellar volumes regressed on cerebral volumes for full data **(a)** and for the 13 species with complete data **(b)** both illustrate isometric scaling trending towards hypo-allometry. Lemuriformes and Hominidea were the two clades with most impressive cerebellar-to-cerebral volume ratios **(Figure 4c)** and were thus specifically colored here (Lemuriformes colored in a bold gray; **a-d**) to highlight their phylogenetic scaling relationships. All species belonging to these infraorders displayed higher cerebellum-to-cerebrum scaling than the primate sample as a whole **(a)**. Zooming in, **(a)** illustrates that while most Hominoidea, including *Homo sapiens, Pongo pygmaeus*, and *Hylobates lar*fall on the regression line of the sample, all but one of the Lemuriformes (*Lemurcatta*; on the line) exceed the general trend. Additionally, in both clades, several species approach isometry, with one member of each falling on the isometric line *(Gorilla gorilla gorilla* and *Mirza coquereli*). **(b)** Illustrates shallowing of the PGLS slope, connected to the smaller sample of volumes. **(c,d)** Ansiform area volume regressed on rest of cerebellar volume **(c)** and cerebral volume **(d)** illustrate hyper-allometric scaling relationships in both cases. Because of the strong positive allometry, species with larger cerebella and cerebra are expected to have larger relative ansiform areas. **(c,d)** Both illustrate that given this observed scaling, Hominoidea do not have unique hyper-scaling of the ansiform area necessarily. Although some Hominoidea lie above this steep regression line, so do several other species (as colored in green; *Mirza coquereli, Aotus trivirgatus, Cebus apella*, and *Papio hamadryas). NS = non-significant*.

#### No distinct allometries in apes

We split the data into ape and non-ape subsets for cerebellum (7 apes (N=24); 27 non-apes (N=39)) and ansiform area (5 apes (N=22); 8 non-apes). We did not find significant interaction with ape membership for cerebellum regressed on cerebrum: F(Df=1, Df_denom_.=30)=1.17, p=.288. Fisher’s R-to-Z transformation indicated significantly worse fit for apes (r=.75, Z=4.37, p<.0001.) and non-apes (r=.87, Z=3.00, p<.01) versus the full model (r=.97). Evidence for difference between groups became even weaker in the 13-species data: F(Df=1, Df_denom_.=9)=.52, p=.49. The full model again better described the data (r=.94) than apes (r=.43, Z= 2.86, p<.01) or non-apes (r=.76, Z=1.70, p=.089) separately. We also found no support in the ansiform area regressions on rest of cerebellar (F(Df=1, Df_denom_.=9)=1.87, p=.204), or cerebral volume (F(Df=1, Df_denom_.=9)=.045, p=.84). The full model outperformed partial models for cerebellar regression (full: r=.85; ape: r=.61, Z=1.19, p=.23; non-ape: r=.90, Z=.53, p=.596) and cerebral regression (full: r=.87; ape: r=.56, Z=1.57, p =.116; non-ape: r=.68, Z=1.14, p=.254). Models fit better in non-apes than in apes for rest of cerebellar (r_apes_=.61, r_non-apes_=. 90, Z=1.72 p=.085) and cerebral regressions (r_apes_=.56, r_non-apes_=.68, Z=.43, p=.667), with only a trend towards significance observed for the cerebellar regression.

Considering this primate-general scaling, we explored if ratio differences within and between clades (Figure 4) could be driven by body size. Linear regressions on body mass^115^ revealed association with ansiform area-to-cerebellum ratios (R^2^_adj_=.44; p<.01) (Supplemental Figure S6a) but not cerebellum-to-cerebrum ratios (R^2^_adj_=-.03; p=.88) (S6b).

### Replication in the Stephan dataset

BM was again the best-supported model (AIC=-19.13) ≈ EB (AIC=-17.51) ≈ OU (diagonal α; AIC=-16.03). It significantly outperformed the star-model (χ^2^=84.60). Pagel’s λ likelihood distributions showed higher likelihoods for higher λ. We adopted BM to map the Stephan 34-species ancestral character estimations (ACEs) to the tree. Cerebellar and cerebral volume ACEs displayed only slightly larger spread and clade-wise distribution was similar. Cerebellum-to-cerebrum ratio evolved similarly as in the main analysis. A notable exception was that Hominoidea had relatively less impressive ratios. Here, relatively large cerebella appeared to be a distinct Strepsirrhini trait (Supplementary Figure S7a). PGLS for cerebellar volume regressed on cerebral volume indicated significant hypo-allometry, with its slope of approximately .92 slightly shallower than in the main analysis (S7b).

## Discussion

### Overview

In the current work, we studied cerebellar volumes for 34 primate species (N=63) and ansiform area volumes for 13 species (N=30). The cerebellum, and the ansiform area specifically, may be part of important cerebello-cerebral loops and networks contributing to associate functions^7,10,13,16,20–23,42,43,50,57,58^. Manual segmentations of the cerebellum and ansiform area were reliable as shown by inter-observer reliability, and comparison with automated^94^ segmentations. The cerebellum scaled isometrically versus the cerebrum in PGLS (coefficient: ~.95). Replication in the Stephan data^84^ argued that the primate cerebellum may in fact scale hypo-allometrically versus the cerebrum (~.92). Conversely, the ansiform area displayed strong positive allometry versus rest of cerebellar (~1.29) and cerebral (~1.25) volumes. Given this scaling across the primate sample, species with larger cerebella and cerebra i.e., hominoids, are expected to have larger relative ansiform areas. Therefore, although we find the largest relative ansiform areas in humans (30.76%) and chimpanzees (27.25%), these ratios are expected from primate-general scaling rules, as illustrated by explorative body mass correlations. We did not find statistical support for distinct allometries in apes.

Together, our findings show that while the cerebellum and cerebrum scale isometrically in primates^50,51,59^, the ansiform area expands relative to the system in a modular and accelerated fashion. Our findings further previous literature, showing that although the ansiform area has become relatively large in humans, this may result from general primate brain architecture and scaling rules.

### Differences between manual and automated segmentations

Manual segmentations were globally similar to automated segmentations in CERES^94^. However, mean manual volumes were systematically higher for the cerebellum. Visual inspection by NM, KH & SLV confirmed accuracy of segmentations. We observed two main differences. First, manual segmentation identified the edges of the cerebellum more exhaustively, whereas automated segmentation was too conservative. Secondly, manual segmentation may have included white matter at the pontine-cerebellar border to separate these structures similarly across all species. Universal guidelines for manual segmentation might help decrease differences, essential for relatively small-sized comparative studies.

### Relative volumes provide advantages

Body mass is strongly coupled to brain size across distant classes such as reptiles, birds^116^, and mammals^117^ including primates^46,69^. The relation between (relative) brain size and intelligence ‘remains one of the thorniest issues in comparative neurobiology’^116^, and simply regressing out body mass may lose important signal. We know that humans have large brains for their bodies^46^. Principal component analysis of cerebello-cerebral phenotypes across mammals separates humans from even primates, a separation that is greatly diminished when body mass is taken out of analysis^118^. We instead believe focusing on reorganization of brain structures themselves may be more revealing for relative importance of brain structures^60,119,120^. Relative volumes provide conceptual and methodological advantages, illustrated by remapping studies that map specific functional areas against a reference area^119,120^. It removes some systematic differences across datasets, species, and specimens, such as acquisition- and shrinkage-related biases (although not tissue-specific shrinkage)^36,84^. Additionally, previously used^70,84,115,121^ compilations report body mass – which is a highly volatile trait – for only a low number of unrelated specimens. Altogether, we did not regress out body mass in analyses.

### The cerebello-cerebral system and Brownian Motion

Our analyses of cerebellar-cerebral system evolution showed strongest support for the Brownian Motion (BM) model. This is in line with previous findings for the evolution of neocortical phenotypes^85^ and with our additional analyses of data from the Stephan collection^84^. High estimates of Pagel’s λ^122^ also supported strong phylogenetic signal. BM argues for trait evolution that is random in extent and direction over time. Such a model of incremental changes, predictable by time since divergence, fits with constrained scaling of the cerebello-cerebral system observed in the current study. At first glance, it clashes with notions of adaptive evolution, as departures from BM have been interpreted to indicate adaptive variation^102,123,124^. The two are not mutually exclusive, as BM models can be supported if selection to specific trait optima is distributed following BM parameters^104^. Recently, generalization of the BM model has been shown to represent both neutral drift and rapid adaptive evolutionary change, the latter of which BM does not do well^125^. It remains open whether homogeneous models (a single regimen per tree, with useful statistical properties^104^) or heterogeneous models (different regimes on subtrees) will best elucidate primate evolutionary complexity^125^.

### The cerebellum and cerebrum scale isometrically

Our study supports primate isometric cerebello-cerebral^50,51,89,118,126^ evolution, tending towards hypo-allometry. A recent study reported similar and significant hypo-allometric cerebello-cerebral scaling across mammals^118^. When we reran PGLS in the Stephan sample, we observed the same cerebello-cerebral scaling coefficient as this study did (.92)^118^. We found no support for distinct scaling patterns in apes.

Lemurs and apes displayed the highest cerebellum-to-cerebrum ratios. Corroboratively, lemurs may have large brains for their bodies^118^, a trait shared with hominoids and especially humans. High relative volumes (Figure 4c) and positive deviations above the slope of the primate dataset (Figure 5a and Supplemental Figure S6b) evidence relatively large lemur and hominoid cerebella. We were unable to detect statistical support for unique ape scaling and did not test lemur uniqueness to prevent hypothesizing after results are known (HARKing)^127^. Although ratios vary markedly across and within clades, median species traits can be closely predicted by general scaling rules. True uniqueness may be found as deviation from this scaling, potentially regressing out appropriately sampled body mass data.

Previous work assessing branch-wise evolutionary rates through the Bayesian reversible jump variable rates model shows that cerebella scale significantly more rapidly than neocortices in great apes^70^. Our approach, examining ancestral characters based on the Brownian Motion model, shows that relatively high cerebellar-to-cerebral scaling may characterize hominoid evolution, with the notable exception of humans. We did not find greater cerebellar scaling versus the greatly expanded neocortex ^89,128^. Accelerated cerebellar volumetric evolution^70^ in great apes may be a response to cerebral expansion. Previous findings of relatively large cerebella – versus what is expected – in contemporary humans and apes, and much less volumetrically pronounced cerebella in recent human ancestors, support this theory^129^, although our data do not. Larger ape samples may shed light on these discrepancies. Alternatively, expansion of the lateral cerebellum may have been a hominoid adaptation^72^, with disproportionate neocortical-to-whole brain expansion^89^ in the lineage leading to humans. A study examining cerebellar^70^ (and ansiform area) evolutionary rates in the current data may be warranted.

### Primate ansiform areas scale hyper-allometrically

The ansiform area scaled to the rest of cerebellum at an approximate 1.3:1 ratio, without unique scaling in apes. This indicates that ansiform area hyperscaling may be a general feature within the examined primates. We offer the cautionary note that we were only able to examine one Strepsirrhini species and could not preclude that the trend may be Haplorhini-specific. Humans and chimpanzees, or hominoids in general, did not differ from allometry more extensively than the supposed lower primates. The strong positive allometry may directly account for high relative ansiform area volumes found in chimpanzees and humans previously^13,130^. Humans may have unique cognitive abilities and exceptional relative brain size^46^, but evolution has not acted to make humans exceptional (the *scale naturæ*). General primate scaling rules logically lead to great trait values and emergent abilities in large-brained humans, supported by increased neuron and connection numbers^131^. Species-specific specializations and reorganizations can occur at any stage of evolution to compound these general changes^132,133^.

Developing structures may become disproportionally large because conserved neurodevelopmental mechanisms scaffold the development of the brain^134^. Small timing differences of cell division cycles may explain the hyperscaling of (neo)cortical surface area in primates^135^ and deviations from isometric cerebello-cerebral scaling observed across mammals^58^ including primates^50,51,53,74^, as well as birds^57^. Primate-general hyperscaling across structures is a common occurrence: predictable relative increases in neocortical volume^89^, white-to-grey matter scaling^136^, cerebellar-to-cerebral neuron numbers^53^, and prefrontal volume^137,138^ have been reported. Hyperscaling of cerebral cortices occurs in simian monkeys^139^ and between macaques and humans^74^ similarly. Truly disproportionate increases within primates may be rare and much of brain organization may be subject to developmental^59^ or functional^65^ constraints. Simultaneously, reorganization has consistently occurred with primate brains generally scaling up – apparently functionally important – connected components. Cognitive modules of the cerebello-cerebral system are primary suspects of such adaptive evolution^7,12,13,140^.

#### Small sample sizes warrant caution

Range of absolute and relative volumes across even the 13-species data was substantial, as expected^80–82^. The ansiform area was observed to consistently take up a disproportionately high percentage of the cerebellum in humans and chimpanzees, fitting well with the 1.3:1 scaling rule. Our data generally resemble prior observations for mean ansiform area volumes in humans (current study: 43.94 cm^3^; Balsters et al.: 53.65 cm^3^; Makris et al.: 43.01 cm^3^), intraspecific variability (5.00 cm^3^; 8.01 cm^3^; 6.38cm^3^), and relative ansiform area-to-cerebellum ratio (30.76%; 36.51%; 29.58%)^13,130^, but illustrate how intraspecific variability cannot be ignored. Reanalysis of ansiform area scaling in larger datasets will be necessary to confirm our observations as they may be partially driven by an underpowered sample and influenced by specimen-specific idiosyncrasies^83,141,142^.

### Community-wide sharing accelerates comparative neurosciences

Reanalysis in the Stephan dataset^84^ offered the following: 1) there is potential for sizable intraspecific samples; 2) but integrating datasets comes with great challenges due to methodological differences, incomplete descriptions, lack of individual observations, or missing provenances; and 3) intraspecific variability is substantial and likely one of the primary drivers of spurious results^80–83,141^. Notable discrepancies in volumetric data between the current study and previous datasets^13,72,84,126,143^ underline the pressing need for intraspecific data in comparative analyses.

Community-wide guidelines for data collection and statistical handling need to be created to facilitate integration of data from primate studies^7,144–146^. Data must be shared openly and completely. Most studies^9,50,70,72,126,143^ contain data provenance but studies describing specimen-specific observations are still rare^72^. Online databases and catalogs for primate brain data – such as PRIMatE Data Exchange (PRIME-DE)^147^ and BrainBox^91^ – can be used to earmark all primate brains with a unique identification number, linked to relevant metadata including sex, age, body weight, captivity status, data availability, and inclusion in specific studies. This would help interpret contradictory results^141^, and test their statistical significance^148,149^. Access to individual observations allows incorporation of intraspecific variability. There is a pressing need to incorporate this statistical uncertainty^80,83^, and statistical methods to do this exist^99,111,121^.

### Summary and outlook

We report cerebellar volumetric data for 34 species (N=63) and ansiform areas (crura I-II) for 13 species (N=30). Our phylogenetic analyses provide corroborating evidence for isometry between cerebellar and cerebral volumes^50,51,118^ and evidence primate-general ansiform area hyperscaling^13,130^. A combination of constrained^59^ and modular^64^ evolution may shape the primate cerebello-cerebral system. The consistent finding of cerebello-cerebral coevolution argues for their structure and function being intimately linked. The strong positive allometry of the ansiform area in primates, potentially alongside that of structurally connected cerebral association cortices^74,75,137,138^, shows detachment from isometric cerebello-cerebral scaling. Exploring associations between ansiform area volume and cognitive ability in relation to intraspecific variability, development, and primate evolution is of prime interest. Larger comparative datasets are necessary: exchange platforms have been initiated. Such datasets in combination with relevant metadata and sizable intraspecific samples will facilitate evolutionary analyses sensitive to intraspecific variability.

The primate cerebello-cerebral system is a strong functional neuroanatomical scaffold^7^. Within it, the ansiform area is characterized by strong positive allometry^13,130^, unique functional activations^14,15^ and involvement in functional networks^16–18,42,43^. It furthermore has a distinct developmental trajectory^150^ and structural properties^150,151^, and a characteristic role in disorders^7^. Concludingly, the ansiform area should be a primary area of interest for the study of human and comparative cognitive correlates in the future^7^.

## Declarations

### Author contributions

**NM.** Conceptualized the manuscript; wrote and adapted custom computer code; performed manual segmentations; performed analyses; wrote the manuscript; incorporated coauthor revisions. **RT.** Provided MRI data; wrote original custom code; wrote used software; provided revision comments manuscript. **VS and MA** provided manual segmentations of ansiform areas for 12 and 6 specimens, respectively. **SBE.** Provided revision comments manuscript; provided funding for the project. **KH*.** Provided MRI data; provided semi-automated segmentation and software support; wrote original custom code; wrote used software; provided revision comments manuscript; provided supervision. **SLV*.** Conceptualized the manuscript; provided a subset of primary segmentations; provided revision comments on manual segmentations, the manuscript, and computer code (multiple occasions); provided supervision; provided funding for the project.

## Acknowledgements

We would like to thank Dr. Carol MacLeod and Dr. Alexandra de Sousa for enthralling discussions about the cerebellum. **RT** and **KH** are supported by the French Agence Nationale de la Recherche, projects NeuroWebLab (ANR-19-DATA-0025) and DMOBE (ANR-21-CE45-0016). **KH** received funding from the European Union’s Horizon 2020 research and innovation program under the Marie Skłodowska -Curie grant agreement No101033485 (Individual Fellowship). Last, this work was funded in part by Helmholtz Association’s Initiative and Networking Fund under the Helmholtz International Lab grant agreement InterLabs-0015, and the Canada First Research Excellence Fund (CFREF Competition 2, 2015–2016), awarded to the Healthy Brains, Healthy Lives initiative at McGill University, through the Helmholtz International BigBrain Analytics and Learning Laboratory (HIBALL), including **NM**, **SBE,** and **SLV**.

## Data availability

Data obtained in the current project, including volumes and ratios, but also intermediate text files and figures, are uploaded to the repository accompanying this project: github.com/NevMagi/34primates-evo-cerebellum. Input files necessary to replicate our results can also be found here, alongside full-size figures to aid legibility. MRI data have been collated from different sources, and published Open Access^85^. The Stephan collection (DOI: 10.1159/000155963) was not published Open Access. Therefore, access to the data was requested by the authors through the Copy Clearance Center Marketplace and facilitated through the Central Library of Forschungszentrum Jülich GmbH. Reuse permission for these data may be requested from the publisher.

## Code availability

Custom computer code was used at all stages of analysis. Some of the scripts used are adapted scripts^85^. Full custom code can be obtained from the GitHub repository accompanying this project: github.com/NevMagi/34primates-evo-cerebellum. Full accreditation of R software used in the project can be found in Supplemental Table S2.

