## Supplemental Methods for "Primate Cerebellar Scaling in Connection to the Cerebrum: A 34-Species Phylogenetic Comparative Analysis"

### Data and provenance

Primate comparative data are rare and sizable datasets are difficult to obtain. We used the collection of primate species collated from different sources in the Brain Catalogue Primates project[1], available on BrainBox[1]. It includes 35 species (N=66) and combines primate MRI data collated from diverse sources, including donations, open data, and new acquisitions (Supplemental File S1). New acquisitions included MRI scans from 31 postmortem specimens (29 species) from the Vertebrate Brain Collection of the National Museum of Natural History in Paris, France[1]. Scans included fully abstracted brains or *in situ* brains, either full body scans or only including the skull, and were acquired on three scanners: two types of 3T Siemens scanners (Tim Trio and Prisma) and an 11.4T Bruker Biospec scanner. For all brains, 3D gradient-echo sequence (FLASH) images were obtained[1]. Beyond these, the Brain Catalogue Primates project[1] includes nine chimpanzee T1-weighted (T1w; 3T) MRI scans from the National Chimpanzee Brain Resource (NCBR; chimpanzeebrain.org)[2], two crab-eating and one rhesus macaque (T1w; 7T) from the Pruszinsky Lab[3], four crab-eating and four rhesus macaques (T1w; 3T) from the Primate Data-Exchange[4], a crab-eating macaque and eastern gorilla downloaded from braincatalogue.org, a bonobo, gibbon, and a western lowland gorilla (T1w; 1.5T) from the NCBR[2], and finally 10 human brains (T1w; 3T) from the ABIDE I dataset[5]. ABIDE I gained approval from the local Institutional Review Boards and had the 18 Health Insurance Portability and Accountability-protected health information identifiers removed[5]. No new primate brains were obtained. Quality control of MRI data including scanning resolutions and signal-to-noise ratios (SNR) can be found in Supplemental File S1[1]. For full data provenance and acquisition, see Heuer et al. (2019)[1]. The 35 species covered the primate phylogenetic tree relatively well, with 9 infraorders coming from both Strepsirhini and Haplorhini suborders. Cebidae (7 species), Hominoidea (7), Lemuriformes (6), and Papionini (5) were most extensively sampled (Figure 1).

### Data preprocessing and quality control

Human[5,6] and non-human primate (NHP)[1] data was already preprocessed. SNRs for the primate specimens can be found in Supplemental File S1.

Comparative datasets vary in data quality, due to differences in brain size, tissue conservation (*in situ* or extracted brains), MRI resolution, unequal SNR, geometrical brain differences, and more idiotypic differences related to specific specimens. During manual segmentation, extensive visual inspection of the data was performed by NM (as well as previously by KH and RT)[1]. Extensive damage beyond the point where it could reasonably be reconstructed was noted in the red howler monkey. This specimen was excluded from the current study. Cerebellar damage was also noted in the black and white ruffed lemur, eastern gorilla, and gibbon. Additionally, two cerebella were not noticeably missing tissue but deformed: the Demidoff's galago and a squirrel monkey. Although these data were included in analyses, their volumetric estimates need to be treated with due diligence. The full damage report can be found in Supplemental File S1.

Lastly, postmortem brains shrink. Comparing postmortem to fresh brain volume[7] is the general practice for quantifying shrinkage. Unfortunately, we could not measure fresh brain volume, nor could we specifically determine brain tissue gravity in our data. We thus did not correct for shrinkage. However, our measures of relative volumes, directly comparing volumes within specimens, should appreciably lessen the effect of shrinkage. Although ideally one would correct individual brain areas by specifically determined shrinkage factors, this is usually not practical and to our knowledge has not been done to do this point[7]. Previous comparisons of volumetric estimates from *in vivo* MRI (Yerkes sample) with postmortem brains (Stephan sample) indicated there was no difference in reliability between datasets[8].

**Manual segmentation procedure**

Manual segmentation is the gold standard of brain segmentation, providing a form of ground truth for brain structure volumes[9]. We obtained initial cerebellar masks semi-automatically, subtracting cerebral masks from whole brain masks using StereotaxicRAMON (github.com/neuroanatomy/StereotaxicRAMON) or using Thresholdmann (github.com/neuroanatomy/thresholdmann) to set thresholds at manually selected locations in the cerebellum directly on the MRI data, and interpolating between these points. Initial cerebellar masks were uploaded to BrainBox[10], allowing the collaborative online segmentation of MRI images. BrainBox visualizes MRI data in the three stereotaxic planes and has several tools for editing and creating segmentations, adjusting contrast settings of the MRI viewer, as well as providing a 3D render of the segmentations. 3D surface reconstructions were only used to assess segmentation quality and not comparative analysis. The authors recognize resolutions of the MRI scans would lead to vast underestimations of cerebellar surface area, which would become more substantial with increasing cerebellar foliation[11,12] and critically depend on resolution. Lastly, we used StereotaxicRAMON for mathematical morphology operations that preserve the original topology of the masks. For an overview of the segmentation procedure, see Figure 2a.

*Cerebellar segmentation*

Manual segmentations (Figure 2b) consisted of brainstem removal, cerebellar surface boundary determination, erroneously marked sulci erasure, and damaged tissue reconstruction. Segmentations were performed by NM, with SLV providing preliminary segmentations for a subset of specimens. Damaged tissue was reconstructed by careful manual interpolation between non-damaged slices. All segmentations were performed using the three stereotaxic planes. First, mid-to-lateral sagittal segmentations were made, after which coronal and axial planes were segmented. All planes were used for subsequent refinement. Because of substantial variation in the quality of brain scans, full grey and white matter volumes were included in segmentations together. Cerebella were cut off from the brainstem by drawing a straight line between adjacent grey matter areas closest to this border, as to prevent biases introduced methodologically based on data quality differences across species. This may have included pontine white matter in cerebellar segmentations. Segmentations were iteratively performed to satisfaction, by alternating mathematical morphology operations and the manual segmentation procedure outlined above. All segmentations (65 cerebella) were visually inspected and approved by two other authors (KH and SLV). See Figure 2b for example cerebellar segmentations in the hamadryas baboon.

*Ansiform area segmentation*

Ansiform area segmentations relied on manual identification of the superior posterior (SPF) and ansoparamedian (APMF) fissures across primates[13–15]. Primate cerebella display conserved but somewhat variable spatial organisation[1,16,17]. Due to its challenging nature and vast differences in data quality, this segmentation was performed in a subset of specimens only. This subset was selected to be as representative of the 34-species phylogenetic tree as possible. However, data quality was highly variable, so 13 species were identified that allowed identification of the fissures with reasonable certainty: 5 apes (N=22 specimens; including 10 humans and 9 chimpanzees) and 8 non-apes. Other data were generally of lower quality, thus raising questions on whether they could be reliably segmented. Hence, and because such segmentations would be extremely laborious, ansiform area segmentations were not made in these data. Humans and chimpanzees – all specimens of appropriate quality – were used to gain an understanding of intraspecies variability of ansiform area volume.

To facilitate accurate segmentation, the SPF and APMF were first identified in the sagittal plane, scanning in mid-to-lateral direction[15]. The area in between these fissures was segmented, not entering vermal portions of lobule VII. As for the whole cerebellum, an iterative segmentation approach in all stereotaxic planes was then adopted. Secondly, an interobserver strategy validated segmentations made by the primary observer (NM). For six species, a secondary observer (MA) provided segmentations alongside NM, both blinded to the other's segmentations. Additionally, a tertiary observer (VS) segmented the remaining human and chimpanzee ansiform areas together with NM in close consultation. All three observers collaboratively trained their identification of the ansiform area on a human brain at 1 $mm^3$ isometric resolution. As a final check, all segmentations (65 cerebellar; 32 ansiform area) were visually inspected by two other authors (KH and SLV). See Figure 2c for an example of ansiform area segmentations in the hamadryas baboon.

After segmentation, a custom python script included in this project's GitHub repository was used through Google Colab to systematically download all data. See Figure 2a-c for an illustration of the full segmentation process.

*Reliability of segmentations*

For the six specimens for which ansiform areas were segmented by both NM and MA, the influence of different observers was quantified through ANOVA as implemented in the *aov* function of R base stats, with the formula:

$$\textit{Ansiform area volume} \sim \textit{species} + \textit{observer} \quad (1)$$

Human cerebellar and ansiform area segmentations were compared to automated segmentations made by CERES[18] at volbrain.upv.es. This was performed similarly through *aov* ANOVA with formulae:

$$\textit{Cerebellar volume} \sim \textit{method} \tag{2}$$

and

$$\textit{Ansiform area volume} \sim \textit{Method} \tag{3}$$

For these comparisons, we additionally calculated the intraclass correlation (ICC) as implemented in the R package irr[19]. An ICC of .75 to .90 is deemed good reliability and a score between .90 and 1.00 indicates excellent reliability[20].

Cerebellar volume was significantly related to the segmentation method (Pr(>F)<.01). ICC was determined at .26 (95% confidence interval (CI): -.38<ICC<.74). Inspection of the volumes obtained through respective methods revealed that manual segmentations were on average 20.4 $cm^3$, or 16.2%, higher than CERES segmentations. Ansiform area volume was not related to method (Pr(>F)=.286), and ICC was .84 (95%CI: .51<ICC<.96). Manual segmentation volumes were only 2.7 $cm^3$ (6.6%) larger on average. Although most volumes were highly similar between both methods, one specimen differed by 6.9 $cm^3$ (18.2%). It represented a rather average-sized human cerebellum, making it unlikely that it significantly altered the results. Full results can be found in the GitHub repository accompanying this project.

Manual ansiform area segmentation volumes were significantly related to species (Pr(>F)<.0001), as expected. However, no significant observer effect was detected (Pr(>F)=.468). Intraclass correlations (ICC) between observers was .986 (95%CI: .92<ICC<1.00). This confirmed that for these six species, the different observers could reliably segment the ansiform area.

**Neuroanatomical measurements**

Neuroanatomical phenotypes of interest included cerebellar, cerebral[1], and ansiform area volumes and cerebellum-to-cerebrum and ansiform area-to-cerebellum ratios. Absolute volumes were measured in $mm^3$ and ratios in percentages. For most analyses, absolute measurements were $\log_{10}$-transformed to facilitate analysis of measurements that differed across multiple orders of magnitude. Volumes were not corrected for shrinkage. Data normality was examined for absolute traits with multiple (>2) observations per species: these included cerebellar and cerebral volumes for four species *(Macaca fascicularis, Macaca mulatta, Homo sapiens,* and *Pan troglodytes troglodytes)* and ansiform area volumes for two species *(Homo sapiens* and *Pan troglodytes troglodytes)* (see Figure 1). Normality was assessed by Shapiro-Wilk tests[21] as implemented in R base stats, and outliers were visualized on boxplots (±1.5 IQR from Q1 and Q3). This led to the exclusion from subsequent analyses of two specimens (one rhesus and one crab-eating macaque; Supplemental Figure S1). Two different crab-eating and rhesus macaque specimens (Supplemental Figure S2) displayed cerebellum-to-cerebrum ratio outliers but were retained in analyses as cerebellar and cerebral volumes fell within the normal range.

**Statistical analyses and reproducibility**

Consensus trees for the 34- and 13-species analyses were obtained from the 10kTrees[22] primate database (10ktrees.nunn-lab.org, version 3). This tree was constructed from 17 genes and 7 different loci[22] (10ktrees.nunn-lab.org/downloads/10kTrees_Documentation.pdf). See Figure 1 for the consensus phylogenetic tree appended with species information, archaeological time scale, and colored by clade.

We used R version R 4.1.0 with the Rstudio version 1.4.1717 graphics user interface for our statistical analyses[23]. Rphylopars[24], which facilitates analysis of multivariate phenotypes for multiple species and multiple individuals per species, was used to test the fit of the extant phenotypes with three of the most common models of trait evolution. We included the neuroanatomical measurements recorded in the current study, as well as the neocortical measures recorded in the same specimens previously[1]. The tested models included the Brownian Motion (BM)[25] model, the Ornstein-Uhlenbeck (OU)[26] model (either with a single alpha value, with an alpha per phenotype, or with a full multivariate matrix of alpha values), and the Early Burst (EB)[27] model. Briefly, BM evolution describes continuously sampled traits varying randomly over time in both direction and extent, leading to trait differences predictable by time since divergence[28]. BM does not simply equate to genetic drift but may also represent weak selection respecting the BM model parameters, or selection towards specific trait optima whose distribution over time can be described by BM[29]. Traits evolving in the OU process behave like BM traits, but with a specific trait optimum, which could be envisioned as a phenotypic value with an adaptive advantage[30,31]. Lastly, EB processes[27] – rapid change after adaptive radiations followed by comparative phenotypic *stasis* – may alternatively explain trait evolution. To assess phylogenetic signal within the data, we chi-squared tested the BM model with lambda fixed at one (data fully explained by phylogeny) versus fixed at zero (star-model, no phylogenetic signal). All models were constructed with the *phylopars* function of Rphylopars[24]. Data fit with the models was determined by the Akaike information criterion (AIC)[24,32], with lower values representing better fit. We regarded a difference between four to seven points to represent significantly more support for the lower-scoring model, whereas a difference exceeding ten represents virtually no support for the model with the higher score[33].

Next, the best-supported evolutionary model was used to reconstruct ancestral states (Ancestral Character Estimation; ACE) for internal nodes of the primate tree – the family structure of the primate dataset – from extant species traits of interest with Rphylopars[24]. In other words, trait estimations were made for every internal node by mapping extant trait values back from their descendants in a way that is consistent with the BM evolutionary model. This implementation is very similar to the ancestral character estimation performed with the ape package[34], and allows incorporation of multiple specimens per species. ACEs were then mapped to the consensus phylogenetic trees for visualization. Phenotypes with observations for all species were mapped to the 34-species tree, whereas those involving the subset of species with ansiform area measurements were mapped to the 13-species tree. For this mapping, ancestral characters were additionally estimated by calculating the BM model with the 13-species phylogeny.

Importantly, extant primate traits cannot be considered to be statistically independent, sharing variable amounts of evolutionary history[28]. Our evolutionary model selection (see Results) indicated a significant phylogenetic signal, which we therefore needed to incorporate in our analyses. The necessity for correction was further illustrated by correlating all neuroanatomical measures in the current study and Heuer et al. (2019)[1], which showed severe collinearity between all variables (Figure 3). Hence, we performed our regressions in the context of phylogenetic generalized least squares (PGLS)[28,35] as implemented in *nlme*[36], which generalizes to phylogenetic independent contrast (PIC) regression under BM[25] forced through the origin[37]. PICs[28] were calculated with ape's[34] *pic.ortho* function, accounting for intraspecific variation. Regressions were performed for four comparisons: cerebellar volume regressed on cerebral volume for all species (1) and for species with complete (including ansiform area) data (2), and ansiform area volume regressed on the rest of the cerebellar volume (3) and cerebral volume (4). Species' median values were used for all regressions.

We furthermore tested whether members of the ape clade displayed different scaling rules versus non-apes. We used ANOVA to test the effect of introducing an interaction term between the independent variable and ape membership in all regressions. We furthermore calculated $R^2_{likelihood}$ values[38,39] for both the full regression models and models with only apes or non-apes included. Consequent Fisher's R-to-Z transformation as implemented in *psych* (cran.r-project.org/web/packages/psych/index.html) allowed us to test whether partial models fit the data better than the model including all primates. Regular non-phylogenetic regressions were also performed but these results were statistically invalid on the assumption that a phylogenetic signal was present in the data. These results are thus not included in the paper but can be found in the accompanying GitHub repository for reference.

Lastly, we reperformed our analysis with data from the Stephan collection[7]. Cerebellar and cerebral volumes were obtained from the publication and matched to primate species available in 10kTrees[22]. This coincidently led to a partially overlapping (in species, not specimens) dataset of 34 primate species. Although comparison between the current and Stephan dataset illustrated the potential for obtaining sizable intraspecific samples, integrating them in a statistically valid manner was currently unfeasible.

doi:10.1007/s12311-022-01390-8.

13. Faber, J. *et al.* Manual sub-segmentation of the cererbellum. 2022.05.09.22274814 Preprint at https://doi.org/10.1101/2022.05.09.22274814 (2022).
14. Larsell, O. & Jansen, J. *The Comparative Anatomy and Histology of the Cerebellum: Vol. 2. From Monotremes through Apes.* vol. 2 (University of Minnesota Press, 1970).
15. Balsters, J. H. *et al.* Evolution of the cerebellar cortex: The selective expansion of prefrontal-projecting cerebellar lobules. *NeuroImage* **49**, 2045–2052 (2010).
16. Sansalone, G. *et al.* Variation in the strength of allometry drives rates of evolution in primate brain shape. *Proceedings of the Royal Society B: Biological Sciences* (2020) doi:https://doi.org/10.1098/rspb.2020.0807.
17. Melchionna, M. *et al.* From Smart Apes to Human Brain Boxes. A Uniquely Derived Brain Shape in Late Hominins Clade. *Frontiers in Earth Science* **8**, 273 (2020).
18. Romero, J. E. *et al.* CERES: A new cerebellum lobule segmentation method. *NeuroImage* **147**, 916–924 (2017).
19. Gamer, M., Lemon, J. & Singh, I. *irr: Various Coefficients of Interrater Reliability and Agreement*. (2010).
20. Koo, T. K. & Li, M. Y. A Guideline of Selecting and Reporting Intraclass Correlation Coefficients for Reliability Research. *Journal of Chiropractic Medicine* **15**, 155–163 (2016).
21. Shapiro, S. S. & Wilk, M. B. An Analysis of Variance Test for Normality (Complete Samples) on JSTOR. *Biometrika* **52**, 591–611 (1965).
22. Arnold, C., Matthews, L. J. & Nunn, C. L. The 10kTrees website: A new online resource for primate phylogeny. *Evolutionary Anthropology* **19**, 114–118 (2010).
23. R Core Team. R: A Language and Environment for Statistical Computing. (2021).
24. Goolsby, E. W., Bruggeman, J. & Ané, C. Rphylopars: fast multivariate phylogenetic comparative methods for missing data and within-species variation. *Methods in Ecology and Evolution* **8**, 22–27 (2017).

25. Freckleton, R. P., Harvey, P. H. & Pagel, M. Phylogenetic analysis and comparative data: a test and review of evidence. *Am Nat* **160**, 712–726 (2002).
26. Uhlenbeck, G. E. & Ornstein, L. S. On the Theory of the Brownian Motion. *Physical Review* **36**, 823–841 (1930).
27. Harmon, L. J. *et al.* Early bursts of body size and shape evolution are rare in comparative data. *Evolution; international journal of organic evolution* **64**, 2385–2396 (2010).
28. Felsenstein, J. Phylogenies and the comparative method. *Am. Nat* **125**, 1–15 (1985).
29. Harmon, L. Phylogenetic Comparative Methods: Learning From Trees. Preprint at https://doi.org/10.32942/OSF.IO/E3XNR (2019).
30. Lande, R. Natural Selection and Random Genetic Drift in Phenotypic Evolution. *Evolution* **30**, 314–334 (1976).
31. Hansen, T. F. Stabilizing Selection and the Comparative Analysis of Adaptation. *Evolution* **51**, 1341–1351 (1997).
32. Akaike, H. A new look at the statistical model identification. *IEEE Transactions on Automatic Control* **19**, 716–723 (1974).
33. Burnham, K. P. & Anderson, D. R. Multimodel inference: Understanding AIC and BIC in model selection. *Sociological Methods and Research* **33**, 261–304 (2004).
34. Paradis, E. & Schliep, K. ape 5.0: an environment for modern phylogenetics and evolutionary analyses in R. *Bioinformatics* **35**, 526–528 (2019).
35. Symonds, M. R. E. & Blomberg, S. P. A primer on phylogenetic generalised least squares. in *Modern Phylogenetic Comparative Methods and their Application in Evolutionary Biology* 105–130 (Springer Berlin Heidelberg, 2014). doi:10.1007/978-3-662-43550-2_5.
36. Pinheiro, J., Bates, D. & R Core Team. nlme: Linear and Nonlinear Mixed Effects Models. (2022).
37. Blomberg, S. P., Lefevre, J. G., Wells, J. A. & Waterhouse, M. Independent Contrasts and PGLS Regression Estimators Are Equivalent. *Systematic Biology* **61**, 382–391 (2012).
38. Ives, A. R. R2s for Correlated Data: Phylogenetic Models, LMMs, and GLMMs. *Systematic Biology*

**68**, 234–251 (2019).

39. Ives, A. & Li, D. rr2: An R package to calculate R2s for regression models. *Journal of Open Source Software* **3**, 1028 (2018).
