## Supplemental Data for "Primate Cerebellar Scaling in Connection to the Cerebrum: A 34-Species Phylogenetic Comparative Analysis"

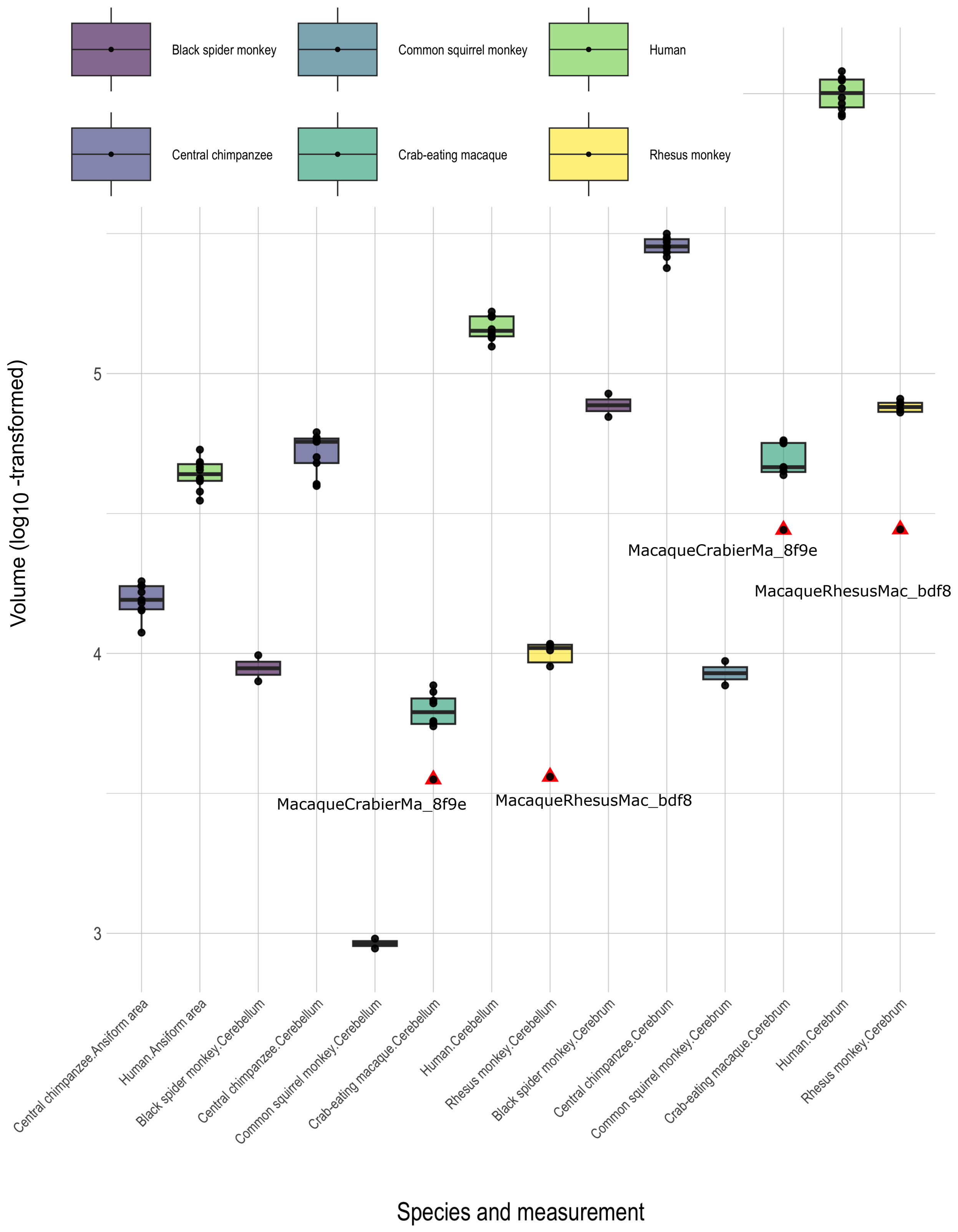


**Figure S1: Outlier detection for absolute volumes.** Box plots illustrate data spread for $log_{10}$-transformed absolute volumes (in $mm^3$) for which multiple specimens were available. Intraspecific ansiform area data were available for humans and chimpanzees only. Two specimens, a crab-eating and rhesus macaque, were identified as outliers based on exceptionally low cerebellar and cerebral volumes.

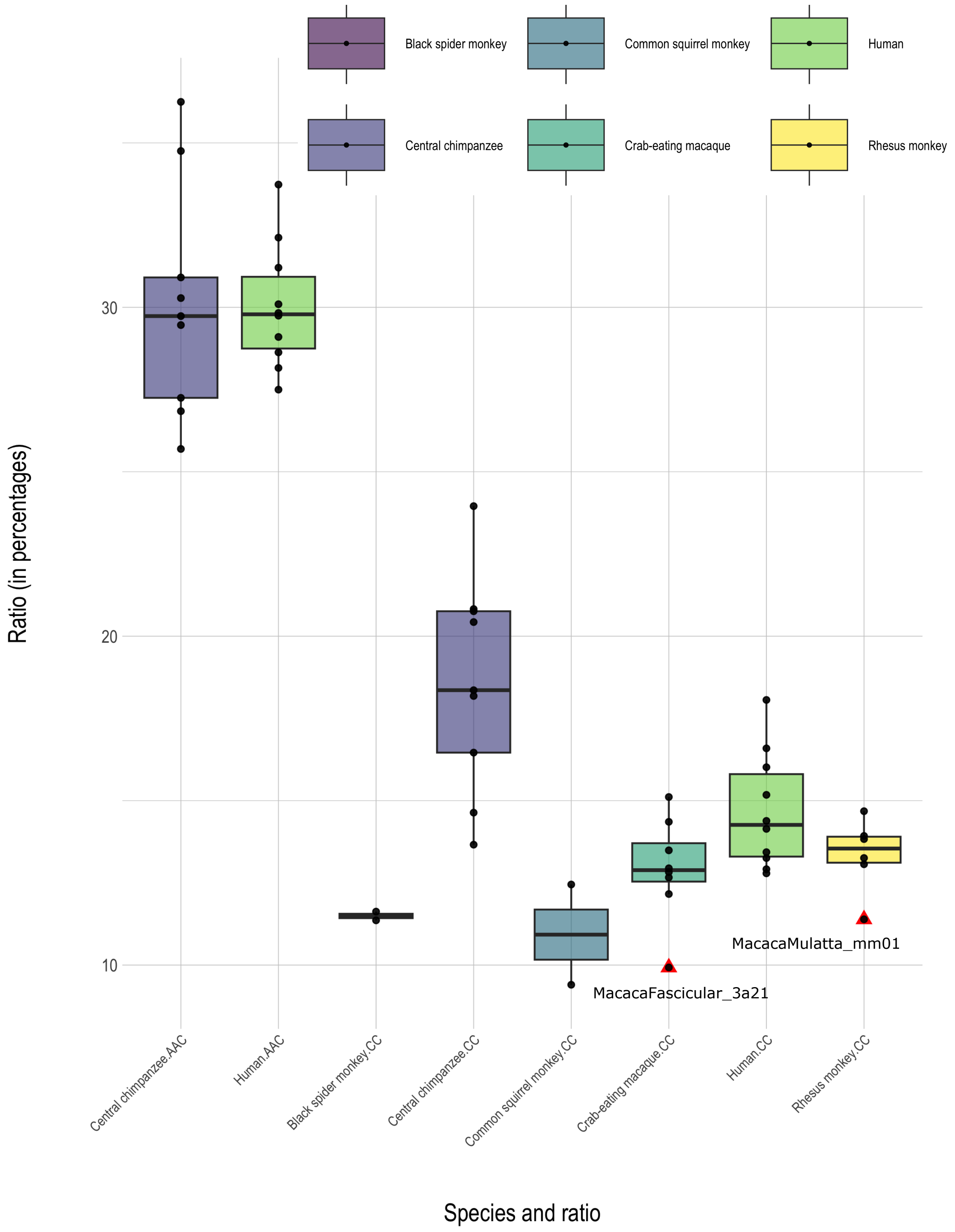


**Figure S2: Outlier detection for volume ratios.** Box plots illustrate data spread for relative volumes (in percentages) for which multiple specimens were available. Intraspecific ansiform area ratios were available for humans and chimpanzees only. Two specimens, a crab-eating and rhesus macaque, were identified as outliers based on exceptionally low cerebellar and cerebral volumes. These individuals were included in consequent analysis, as their absolute values occupied a position within the normal range. AAC = ansiform area-to-cerebellar volume ratio; CC = cerebellar-to-cerebral volume ratio.

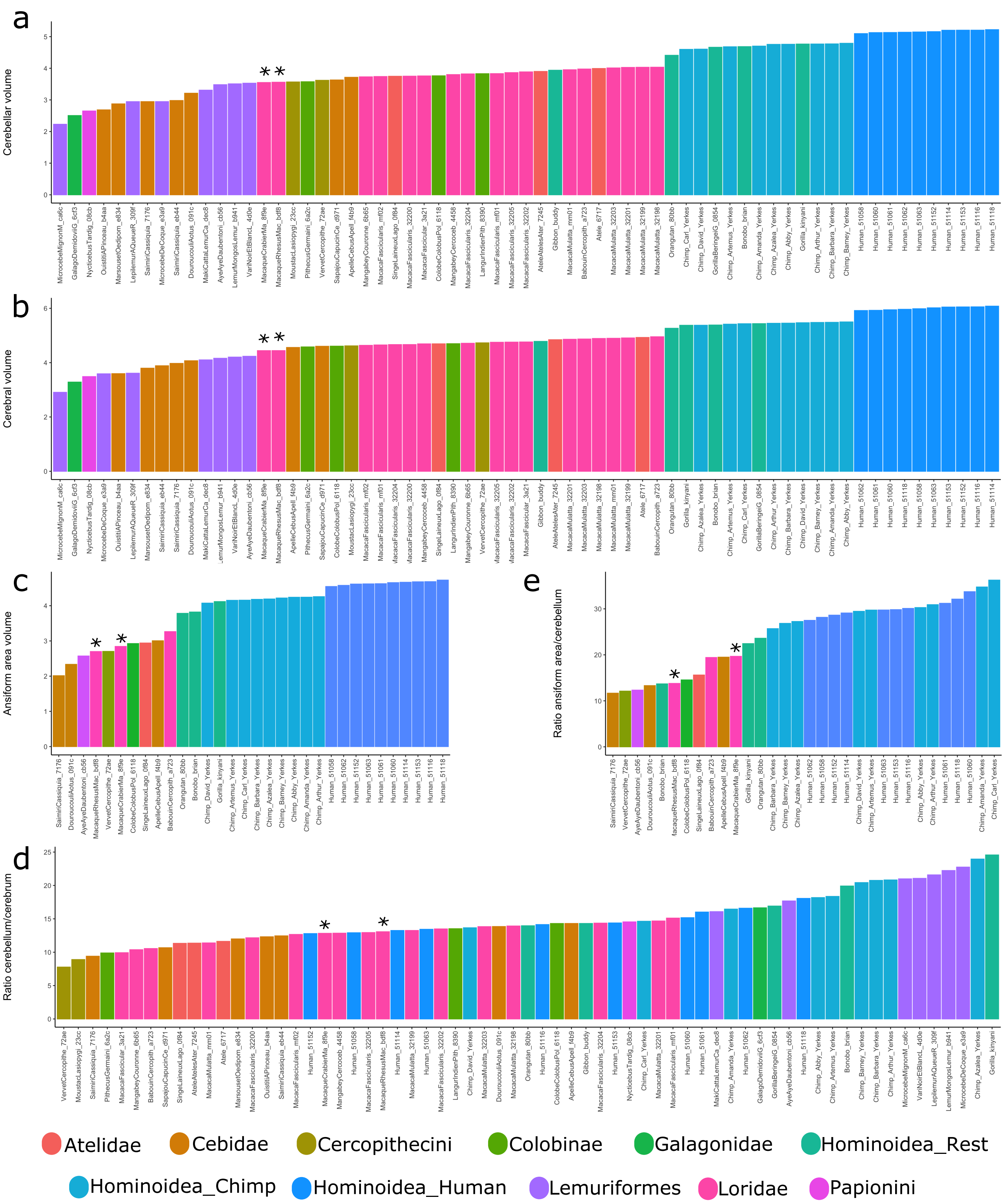


**Figure S3: Absolute and relative traits per clade.** Traits are ordered based on size and colored by clade. Hominoidea are split into humans, chimpanzees, and remaining species (see legend). Absolute traits **(a-c)** (in $mm^3$, $log_{10}$-transformed) were systematically largest in apes and specifically humans, whereas ordinal relative traits **(c-d)** (in percentages) were spread more across clades. Full specimen IDs are provided on the x-axis. Two specimens that were excluded are marked by asterisks in all subfigures.

**Table S1: Ancestral character estimations (ACEs) for absolute and relative neuroanatomical measurements.** ACEs are provided for cerebellar, cerebral, and ansiform area volumes (in $mm^3$), as well as cerebellum-to-cerebrum and ansiform area-to-cerebellum ratios (in percentages). ACEs incorporated intraspecies variation when multiple observations were available **(main text, Table 1)** and were constructed from the variance-covariance matrix derived from the 34-species consensus phylogenetic tree **(main text, Figure 1)**. Estimations for absolute values are rounded to the nearest integer, and estimations for percentages to the second decimal. Full, unrounded estimations can be found in the GitHub repository accompanying this publication. Internal node numbers refer to nodes in the 34-species tree **(main text, Figure 1)**.

| *Internal node* | *Cerebellar volume* | *Cerebral volume* | *Ansiform area volume* | *Cerebellum-cerebrum ratio* | *Ansiform area-cerebellum ratio* |
|---|---|---|---|---|---|
| 35 | 1,856 | 11,968 | 438 | 16.02 | 15.74 |
| 36 | 3,686 | 26,789 | 724 | 14.26 | 17.05 |
| 37 | 7,928 | 58,707 | 1,516 | 14.02 | 19.01 |
| 38 | 6,361 | 52,050 | 1,101 | 12.61 | 17.92 |
| 39 | 5,966 | 53,030 | 1,072 | 11.51 | 17.91 |
| 40 | 4,864 | 48,373 | 804 | 10.19 | 16.83 |
| 41 | 6,349 | 55,347 | 1,194 | 11.70 | 18.34 |
| 42 | 6,411 | 56,221 | 1,249 | 11.59 | 18.52 |
| 43 | 7,183 | 62,380 | 1,453 | 11.62 | 18.96 |
| 44 | 7,680 | 60,170 | 1,432 | 13.00 | 19.08 |
| 45 | 5,787 | 47,060 | 902 | 12.64 | 17.17 |
| 46 | 5,513 | 45,618 | 799 | 12.41 | 16.70 |
| 47 | 16,614 | 110,301 | 3,529 | 15.57 | 21.55 |
| 48 | 26,174 | 163,928 | 5,944 | 16.43 | 23.33 |
| 49 | 48,447 | 282,748 | 11,944 | 17.41 | 25.85 |
| 50 | 50,559 | 259,653 | 12,317 | 19.44 | 25.13 |
| 51 | 60,243 | 360,494 | 15,400 | 16.96 | 27.11 |
| 52 | 50,191 | 285,005 | 11,645 | 17.88 | 25.57 |
| 53 | 2,251 | 17,856 | 385 | 12.92 | 15.33 |
| 54 | 5,320 | 43,875 | 882 | 12.61 | 17.00 |
| 55 | 2,015 | 15,991 | 344 | 12.89 | 15.08 |
| 56 | 1,894 | 14,960 | 327 | 12.94 | 15.02 |
| 57 | 1,324 | 10,433 | 258 | 13.00 | 14.83 |
| 58 | 2,055 | 16,495 | 339 | 12.70 | 14.92 |
| 59 | 4,060 | 31,774 | 659 | 13.02 | 16.67 |
| 60 | 1,418 | 8,727 | 360 | 16.70 | 15.23 |
| 61 | 1,483 | 8,592 | 382 | 17.72 | 15.12 |
| 62 | 1,239 | 6,518 | 423 | 19.35 | 15.90 |
| 63 | 976 | 5,014 | 412 | 19.76 | 16.40 |
| 64 | 513 | 2,588 | 344 | 20.20 | 16.97 |
| 65 | 2,047 | 10,678 | 485 | 19.42 | 15.15 |
| 66 | 2,242 | 11,874 | 459 | 19.15 | 14.57 |
| 67 | 674 | 4,215 | 197 | 16.20 | 14.22 |

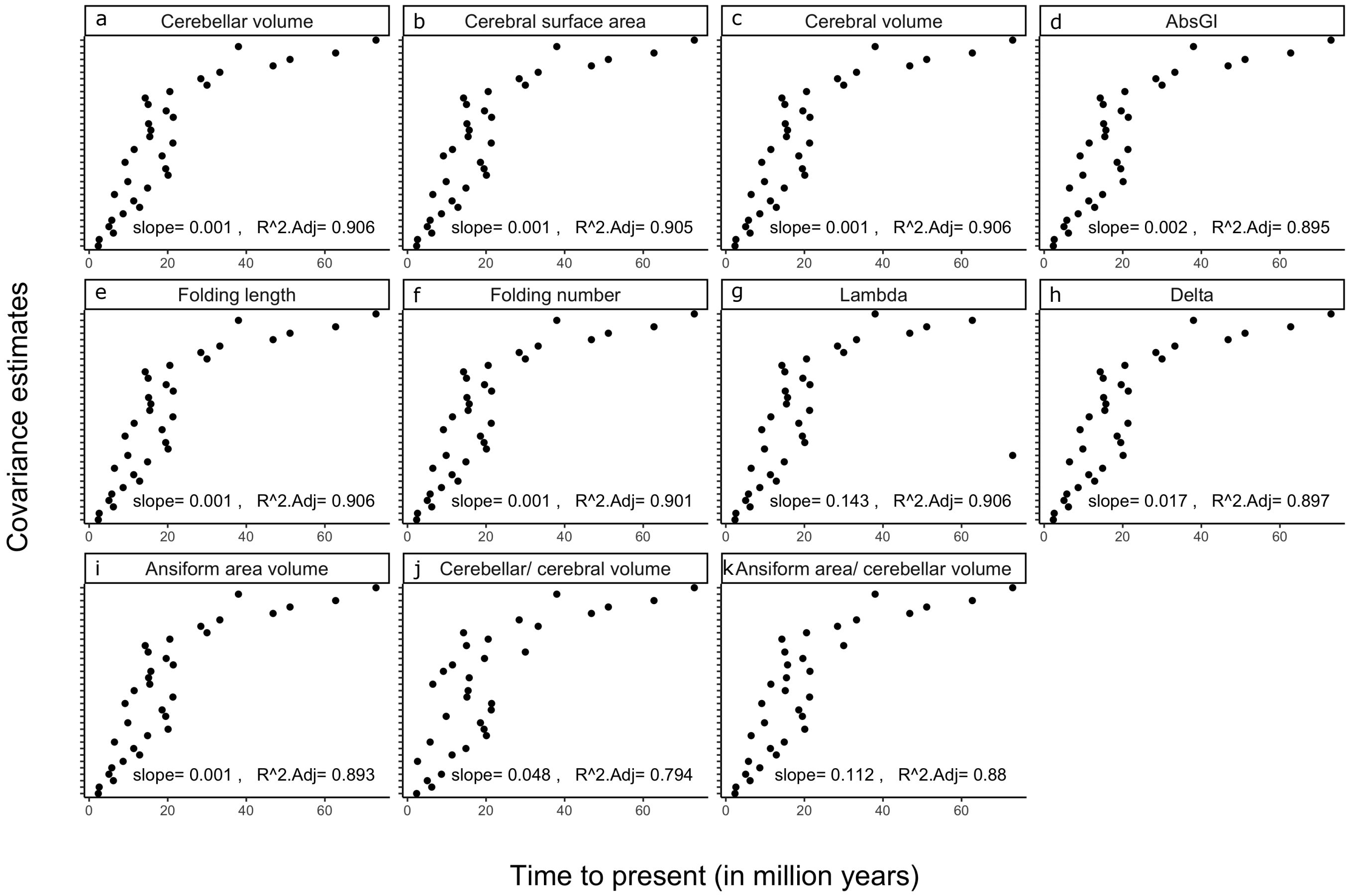


**Figure S4: Uncertainty at ancestral nodes.** To explore the uncertainty of ancestral character estimations (ACEs) at ancestral nodes, covariance estimates at each internal node were correlated with time to present. For all measures **(a-k)**, including those from **Heuer et al. (2019) (b-h)**, covariance correlated strongly with evolutionary time. In the measurements reported in the current study **(a,b,i-k)**, ACEs were thus more uncertain for early nodes than for more recent ones.

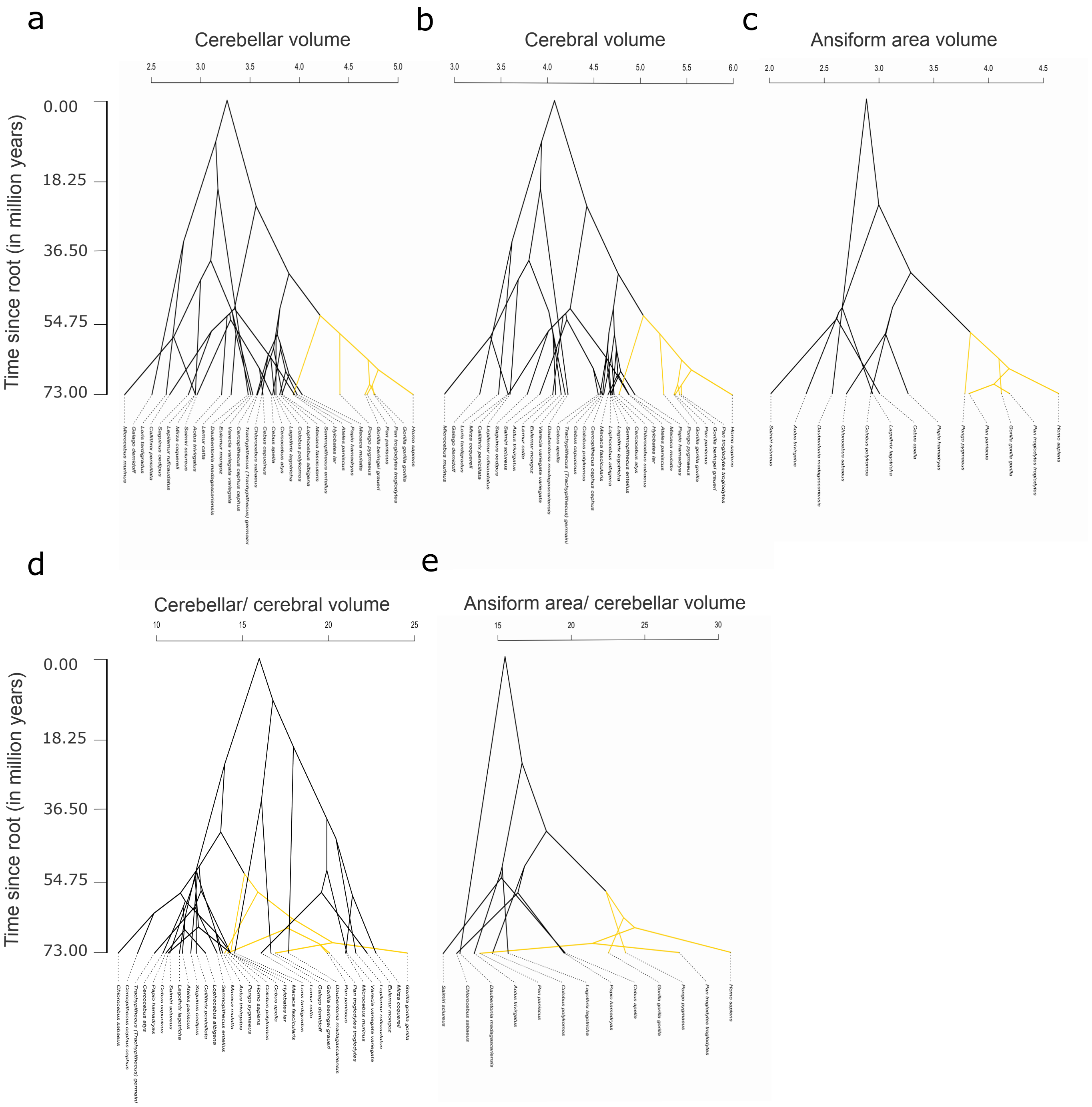


**Figure S5: Phenograms for absolute and relative traits**. Alternative visualization of the traits in extant species and ancestral character estimations (ACEs), that emphasizes trait values specifically. The ape-clade is colored yellow. Absolute volumes (in $mm^3$, $log_{10}$-transformed) (**a-c**) separated apes from non-apes, with *Hylobates lar* (common gibbon) as exception. Cerebellum-to-cerebrum ratios (in percentages) (**d**) were more mixed between apes and non-apes. Except for *Pan paniscus* (bonobo), ratios of ansiform area-to-cerebellum (**e**) strongly separated apes from non-apes.

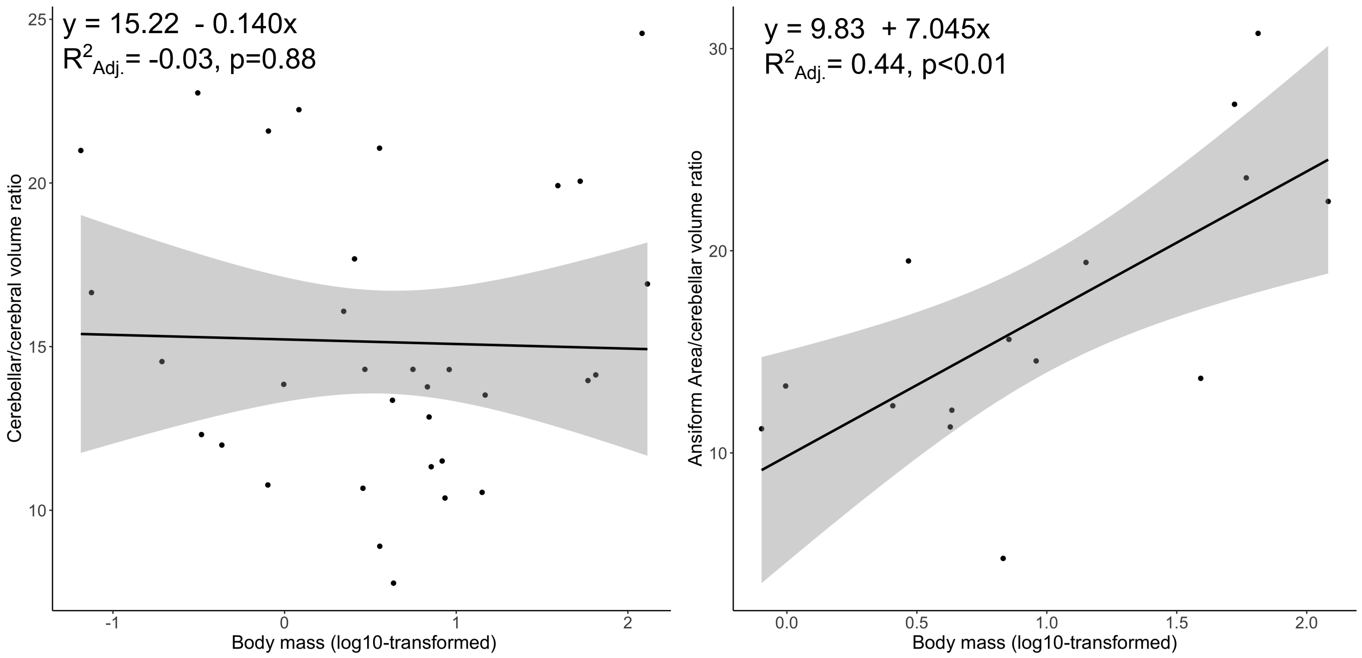


**Figure S6: Explorative regression of volumetric ratios on body mass.** We explored whether cerebellum-to-cerebrum **(a)** and ansiform area-to-cerebellum **(b)** ratios could be directly related to $\log_{10}$-transformed body mass. Linear regression showed no detectable relationship between relative cerebellar volume and body mass **(a)**. However, relative ansiform area volume **(b)** showed association with body mass ($R^2_{Adj.}$ = .44, p<.01). This indicated that relative ansiform area volumetric variability was partly driven by body mass, with larger species having higher ratios. Body mass is given in kilograms. Adj. = adjusted.

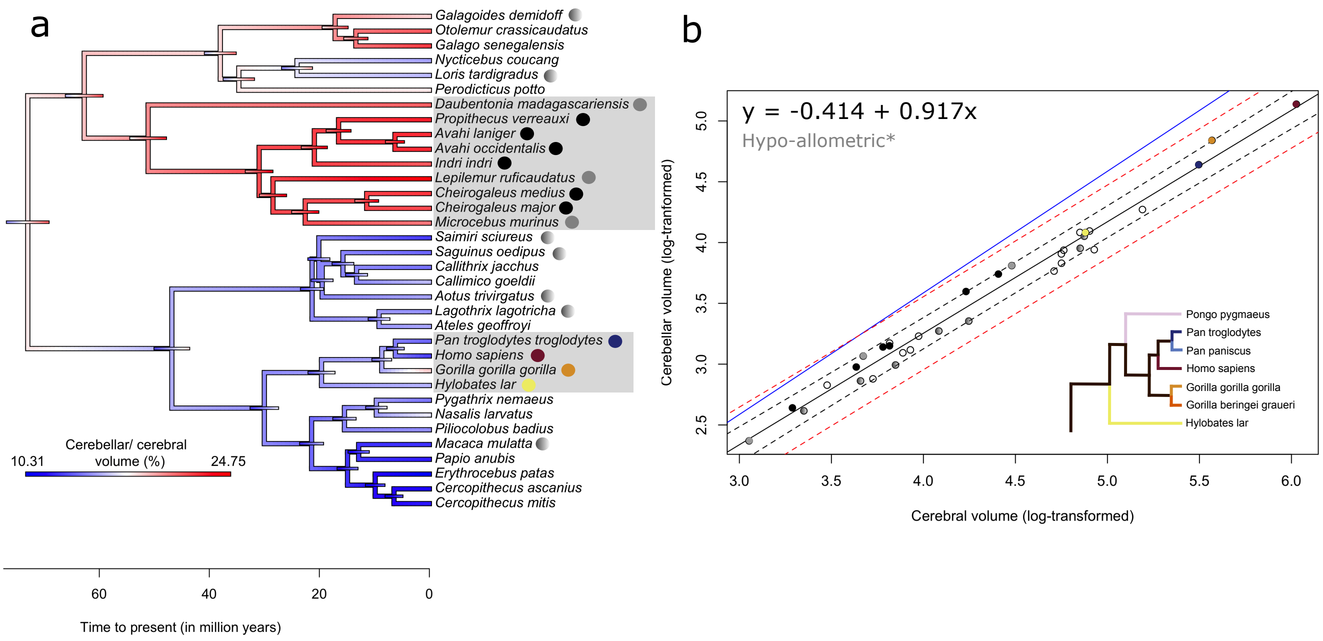


**Figure S7: Replication of allometric relationship between the cerebellum and cerebrum in the Stephan collection.** The **Stephan et al. (1981)** collection reported cerebellar and cerebral volumes for 34 species available in 10kTrees **Arnold et al. 2010**. Ancestral character estimations (ACEs) **(a)** and phylogenetic generalized squares regression (PGLS) **(b)** were repeated in this complementary data set. Despite only partial overlap in species, ACEs for cerebellar-to-cerebral volume ratios **(a)** largely mirrored the main analysis **(main text, Figure 4c)**, with quantitatively similar distributions of ratios within the Platyrrhini and the wider Strepsirrhini clades. In this dataset however, Hominoidea displayed some of the lower ratios, with only *Gorilla gorilla gorilla* (Western lowland gorilla) having relatively high ratios. In contrast to the current study, these data indicated that large relative cerebella were a distinct Strepsirrhini trait. Like in the main analysis, Lemuriformes displayed consistently high ratios. PGLS regression between cerebellar and cerebral volumes in the Stephan collection **(b)** illustrated qualitatively similar scaling as the primary analysis **(main text, Figure 5a)**. However, the slope of the allometric relationship was shallower at approximately .92, and its 95% confidence intervals excluded the isometric line. Therefore, the Stephan collection data revealed significant hypo-allometric scaling of the cerebellum to the cerebral cortex. All but one of the Lemuriformes (*Daubentonia Madagascariensis*; on the regression line) have values above the regression line, approaching isometry and mirroring the main analysis **(main text, Figure 5a)**. In Hominoidea, *Homo sapiens* and *Hylobates lar* fall on the regression line (as in the main analysis), as does *Pan troglodytes troglodytes* (which is hyper-allometric in the main analysis). Reflecting the high ratio of cerebellar-to-cerebral volume as seen in **(a)**, *Gorilla gorilla gorilla* is the only Hominoidea species above the regression line (**b**), approaching isometry (it scales isometrically in the main analysis). Legend: apes are colored as in **(main text, Figure 5a)**, with legend again shown here in **(b)**. Lemuriformes are colored here as well, with solid greys representing the species overlapping with the main analysis, and solid blacks indicating species unique to the Stephan dataset **(a,b)**. Lastly, the other species overlapping between both data sets, that were not specifically considered, are colored a shaded grey **(a,b)**.

**Table S2: R software packages used in this study.** Software used for analyses or data handling in this study that are not included in base R are listed. Authors of packages or functions are accredited under source.

| **Package or *function*** | Source |
|---|---|
| ape | Paradis, E & Schliep, K (2019). ape 5.0: an environment for modern phylogenetics and evolutionary analyses in R. Bioinformatics 35, 526–528. |
| corrplot | Wei T & Simko V (2021). R package 'corrplot': Visualization of a Correlation Matrix. (Version 0.92), https://github.com/taiyun/corrplot. |
| data.table | cran.r-project.org/web/packages/data.table/index.html. |
| dispRity | Guillerme T (2018). "dispRity: A modular R package for measuring disparity." *Methods in Ecology and Evolution*, **9**(7), 1755-1763. |
| evomap | Smaers, JB: github.com/JeroenSmaers/evomap. |
| ggpattern | github.com/coolbutuseless/ggpattern. |
| ggplot2 | Wickham H (2016). ggplot2: Elegant Graphics for Data Analysis. Springer-Verlag New York. ISBN 978-3-319-24277-4, https://ggplot2.tidyverse.org. |
| *ggplotRegression* | Johnston, S: sejohnston.com/2012/08/09/a-quick-and-easy-function-to-plot-lm-results-in-r/. |
| ggpmisc | Aphalo, PJ (2016) Learn R ...as you learnt your mother tongue. Leanpub, Helsinki. |
| lsmeans | Lenth, RV (2016). "Least-Squares Means: The R Package lsmeans." Journal of Statistical Software, 69(1), 1–33. doi:10.18637/jss.v069.i01. |
| nlme | Pinheiro J, Bates D, R Core Team (2022). nlme: Linear and Nonlinear Mixed Effects Models. R package version 3.1-161, https://CRAN.R-project.org/package=nlme. |
| nortest | https://cran.r-project.org/web/packages/nortest/index.html. |
| phytools | Revell LJ (2012). "phytools: An R package for phylogenetic comparative biology (and other things)." Methods in Ecology and Evolution, 3, 217-223. |
| psych | Revelle, W (2022). psych: Procedures for Psychological, Psychometric, and Personality Research. Northwestern University, Evanston, Illinois. R package version 2.2.9, https://CRAN.R-project.org/package=psych. |
| Rphylopars | Goolsby, E. W., Bruggeman, J. & Ané, C. Rphylopars: fast multivariate phylogenetic comparative methods for missing data and within-species variation. Methods in Ecology and Evolution 8, 22–27 (2017). |
| rr2 | Ives et al., (2018). rr2: An R package to calculate $R^2$s for regression models. Journal of Open Source Software, 3(30), 1028, https://doi.org/10.21105/joss.01028. |
| tidyverse | Wickham et al., (2019). Welcome to the Tidyverse. Journal of Open Source Software, 4(43), 1686, https://doi.org/10.21105/joss.01686. |
